# Multi-area Decision Dynamics Across Human Cortex Shape Confidence

**DOI:** 10.1101/2025.10.28.684827

**Authors:** Alessandro Toso, Ayelet Arazi, Jaime de la Rocha, Konstantinos Tsetsos, Tobias H. Donner

**Affiliations:** Department of Neurophysiology and Pathophysiology, University Medical Center Hamburg-Eppendorf, 20246 Hamburg, Germany; Institut d’Investigacions Biomèdiques August Pi i Sunyer, Barcelona, Spain; Trinity College Institute of Neuroscience and School of Psychology, Trinity College Dublin, Dublin, Ireland; Bernstein Center for Computational Neuroscience, Berlin

## Abstract

Theories of the neural basis of decision-making postulate a competition between choice-selective populations of neurons. Mounting evidence shows that choice-selective neural activity is distributed across many brain areas. We hypothesized that these distributed choice codes shape agents’ confidence in their decision. To test this idea, we tracked the distributed dynamics of candidate neural decision variables via magnetoencephalography while human participants reported their decisions and the associated confidence in a challenging visual evidence accumulation task. Premotor and motor cortex showed the strongest and earliest choice-selective activity across the cortex. Premotor signals supporting the selected and non-selected choice options reflected a weak, intrinsic competition, and both made unique contributions to confidence of opposite sign. Confidence also depended on choice codes beyond those expressed in premotor and motor cortex, in clusters of areas in prefrontal and parietal association cortex. Also here, regions supporting the selected and non-selected options uniquely contributed to confidence with opposite sign. We conclude that the sense of confidence emerges from decision-related signals that are distributed across many cortical areas.

## Introduction

Mammals, and humans in particular, are equipped with a well-calibrated sense of confidence, the belief about the accuracy of their decisions^1,2^. A substantial body of work has illuminated the psychophysical properties of this important cognitive ability and delineated its precise functions in the online control of behavior, including the regulation of speed-accuracy tradeoffs ^3^, history-dependent biases ^4,5^, learning ^6^, exploration and sampling ^7^, and decision-making in volatile environments mixing different forms of uncertainty^8,9^.

The temporal integration of sensory evidence about the state of the environment is a core computation in dynamic models of decision and confidence formation^10–12^. A prominent such model, the drift diffusion model, describes the dynamics of a single decision variable, which integrates fluctuating decision evidence towards one of two bounds^10,13,14^, and accounts for choice and reaction time (RT) across a wide range of tasks^10,13,15^. One-dimensional accumulator models like drift diffusion need additional mechanisms, such as collapsing bounds or a learned mapping between decision time and accuracy, to account for trial-to-trial variation in confidence^16,17^. Models entailing a race between different accumulators for each choice option provide access to the accumulated evidence supporting also the losing option^18^. Such models propose that confidence depends only on the value of the losing decision variables at the time of commitment, which takes different values from trial to trial^19,20^. Yet another class of models postulates that decisions result from multiple loosely coupled “decision modules” each implementing their own race between accumulators; confidence depends on the number of modules supporting the selected choice, akin to a vote-counting mechanism^21–23^. Key experimental data to evaluate these different architectures have been largely missing, because the simultaneous tracking of many choice-selective neural signals distributed across the cortex has been challenging.

In decision-related areas of the cerebral cortex, groups of neurons are selective for different choice options and ramp up during decision formation^10,18,24–26^. While such choice-selective activity is strong in motor and premotor cortex in humans^11,27–29^, monkeys^30,31^, and mice^32^, it also occurs in several regions of association cortex^12,33–36^. We hypothesized that the collective dynamics of all these choice-selective neural signals, so-called neural decision variables (DVs)^30^, shapes confidence. Neural circuit models postulate that different neural DVs are coupled via inhibitory interactions within areas^18,26,37^ and through recurrently excitatory interactions between areas^34,38^. We reasoned that such inhibitory crosstalk may generate intrinsic (stimulus-independent) correlations between neural DVs. Therefore, pinpointing the unique contribution of these distributed and competing neural DVs to confidence requires tracking multiple DVs concurrently during a single decision. Recent advances combining magnetoencephalography (MEG) source imaging^39^ and regionally-specific decoding techniques have opened up a window on distributed cognitive dynamics in the human cortex^11,27,40,41^. This combination affords tracking multiple neural DVs with sufficient temporal resolution to track the dynamics of neural DVs, sufficient spatial resolution to delineate multiple regional DVs, and wide spatial coverage to measure DVs concurrently across many cortical regions^11^.

Building on these advances, we here developed an approach to assess the interactions between competing neural DVs and quantify their unique impact on decision confidence. We combined a visual evidence accumulation task with behavioral modeling and multi-variate decoding analyses of human MEG data. The task required the magnitude comparison of two independently fluctuating input streams and was tailored to identifying statistical signatures of intracortical competition in the neural DVs. These signatures were identified for the proxies of the two competing neural races within premotor (and motor) cortex, where choice-predictive activity was stronger. Both the “winning” and the “losing” neural DVs in this part of cortex made a unique statistical contribution to trial-by-trial variations in confidence, with a stronger impact for the winning DV. Critically, also DVs in association cortex made a significant contribution to confidence, over and above the (pre)motor neural DVs: confidence increased with the mean strength across the winning neural DVs in posterior parietal and prefrontal cortical areas, and confidence decreased with the mean strength across the losing DVs in these areas. Our study opens a new route for relating distributed cortical computations to subjective mental states such as confidence.

## Results

During MEG recordings, twenty participants compared the strength of two visual stimuli that were presented simultaneously in the left and right visual hemifields (Figure 1A; see Methods). Each stimulus consisted of a sequence of ten successive circular gratings (100 ms duration), which fluctuated in contrast. The contrasts of each grating provided the input samples, which were randomly and independently drawn from either a high-contrast or a low-contrast distribution (Figure S1A, B). In each trial, one of these two distributions was randomly assigned to the left and one to the right stimulus location, defining two stimulus categories, Left and Right. High and low contrast distributions were Gaussians with a common standard deviation and a mean that differed by a value, delta-contrast, that determined the “evidence strength” governing the difficulty of the decision. Different from commonly used evidence accumulation tasks^10,27,42,43^, the fluctuations of the evidence (i.e., contrast samples) around the mean were independent for the left and right sides (Figure S1, C,D) Participants were asked to infer the stimulus category and report it by a button press with the left or the right hand. For each hand, they also selected one of two possible buttons to report their confidence (high versus low) associated with the decision. This joint choice and confidence report was prompted by the stimulus offset. RT was measured as the time from stimulus offset to button press. At the end of the trial, participants received auditory feedback about the correctness of their decision.

**Figure 1.**
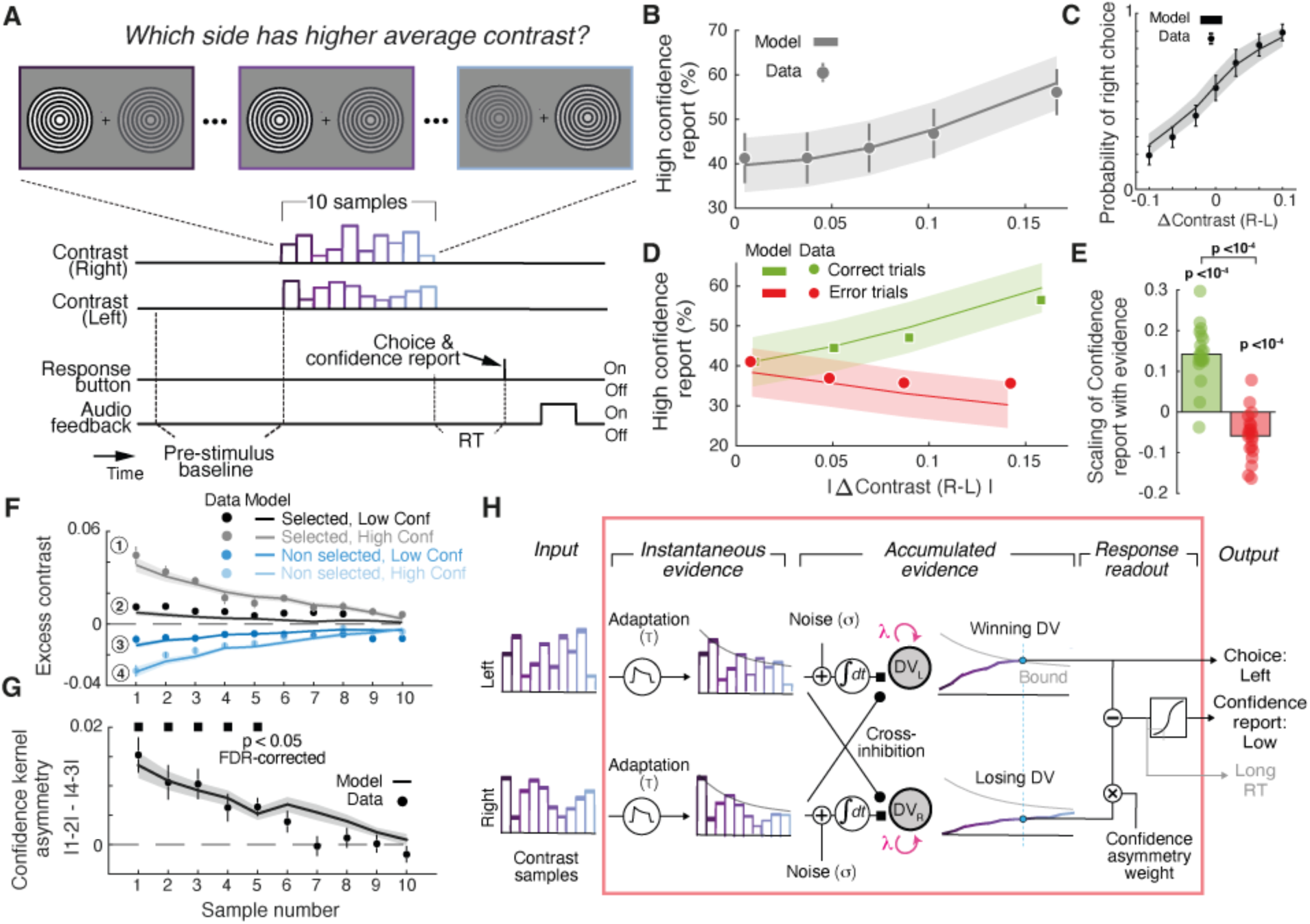
Behavioral task and confidence signatures. **(A)** Schematic of events during an example trial. A sequence of fluctuating contrast samples is shown in each visual hemifield. After stimulus offset, participants report the side with the larger mean contrast (left versus right) combined with their confidence in the correctness of that judgment (high versus low) by pushing one of four buttons. **(B)** Probability of high confidence as a function of evidence strength (absolute value of delta contrast). **(C)** Probability of right-side choice as a function of average delta contrast between right and left stimuli. Data points, group average; vertical lines, SEM across participants; shaded area, SEM of model predictions. **(D)** Probability of high confidence report as a function of absolute value of delta contrast, split by correct and error trials. **(E)** Correlation between high confidence report with evidence strength for correct (green) and error (red) trials. Data points, individual participants; bars, group average. P-values obtained from two-sided permutation test against 0. **(F)** Excess contrast fluctuations sorted by the side selected by the participant and the associated confidence report. **(G)** Asymmetry of confidence kernels, computed as the difference between confidence kernels for selected (1-2) and non-selected (4-3) side in panel F. Black horizontal squares indicate p < 0.05 (two-sided permutation-test against 0, FDR-corrected). All panels except C,E: Data points, group average; vertical lines, SEM across participants; lines, average predictions of a competing accumulator model (Methods); shaded area, SEM of model predictions. **(H)** Model schematic. The probability of “high confidence” report is computed as the difference between the winning and a weighted (weights between 0 and 1) version of the losing DV, transformed through a logistic function. The leak parameterizes drift in the memory: λ = 0 implies perfect integration; λ > 0 indicates leaky (forgetful) integration; λ < 0 indicates unstable integration. See main text for details. Data from N = 20 participants.

Although a staircase procedure kept the evidence strength in a narrow range around 75% correct (Methods), choice probability lawfully scaled with its signed value (Figure 1C). Trial-to-trial fluctuations of the input samples around the mean also affected choice (Figure S1E and Methods). This impact, captured by the so-called psychophysical kernel^11,44–46^, was significant for all ten samples in the stimulus sequence, but decayed across time in the majority (18 out of 20) individuals (Figure S1E,F). This indicates that participants used all samples to compute their choices, in line with evidence integration, but most of them systematically deviated from uniform integration in that they systematically overweighted earlier samples compared to later ones (“primacy”) as observed previously in monkeys and humans^11,45^.

### Behavioral signatures of decision confidence

The probability of high confidence reports increased with evidence strength (Figure 1B), specifically for correct trials, combined with a decrease with evidence strength for error trials (Figure 1D, E). Further, fluctuations on the selected side had stronger impact on confidence reports than fluctuations on the non-selected side (Figure 1F, G). To measure this asymmetry, we calculated the “excess contrast” for the selected and non-selected side as the deviation of each sample from the respective generative mean (Methods). There was an average excess of evidence on the chosen side, and a negative excess in the non-chosen side (Figure 1F). When splitting by reported confidence, both the excess and lack of evidence were more pronounced in high compared with low confidence trials (Figure 1F). This is expected as larger excess on the selected side and larger lack of evidence in the non-selected side both result in stronger evidence for the selected side and, correspondingly, greater confidence in that choice (Figure 1B). Critically, this effect was asymmetric: fluctuations on the selected side had a stronger impact on confidence than fluctuations on the non-selected side (Figure 1F,G). Such asymmetry can be explained by a differential weighting of the input for choice and confidence reports^44^ or by a single inference underlying both judgments, but only using part of the input sequence (e.g. due to a collapsing bound; see below)^17^. In sum, participants were able to provide well-calibrated confidence reports that were consistent with statistical decision theory^17,47^ as well as previous behavioral findings from similar tasks^4,44,48^.

The behavioral signatures of confidence were all captured by an accumulator model of decision formation (lines in Figures 1 and Figure S3 are model fits). The model consisted of three processing stages (Figure 1H; Methods): (i) a sensory module transforming the two stimulus streams into two streams of instantaneous decision evidence through an adaptation kernel^11,14,49^; (ii) an accumulator module integrating the two instantaneous evidence streams into the two DVs, with cross-inhibition between the two streams, up to an absorbing and collapsing bound; (iii) a readout module transforming the two DVs into behavioral choices and confidence reports. The readout module determined choice as the DV first crossing the bound, or the larger DV if no bound was crossed. Confidence was computed as a transformation of the difference between winning and losing DV at the decision time, whereby the losing DV was scaled by a so-called “confidence asymmetry weight”. The model had 11 free parameters, which were fitted to reproduce the choices and confidence reports of each participant (Methods). Despite the large number of parameters, the model provided better cross-validated fits to the data than several alternative models (Figure S2A; Methods), and it was well recoverable (Figure S2B). The model also reproduced the observed psychophysical kernels with smaller deviation (mean-squared error) than all other models considered (Figure S2C) as well as several key signatures of the neural dynamics, as we will show in subsequent sections.

The above signatures of confidence reports were mirrored in the measured RTs and the model’s decision times (Figure S3). Participants’ RTs and confidence judgments were weakly but significantly correlated (group mean of single-trial correlation coefficient: -0.162; p < 10^-4^; Figure S3A). While the time of commitment is not well identified in our current fixed duration task (see Discussion), we estimated model decision times for each trial as the mean time of bound crossing across many simulations (different instances of noise) for that trial’s evidence sequence (Methods). Also the RTs showed the crossover pattern as function of evidence for correct and error trials (Figure S3B-D) and the asymmetry of psychophysical kernels for selected versus non-selected sides (median split based on RT; Figure S3E-F). In other words, RTs exhibited the same signatures of decision confidence as the confidence reports, just with flipped sign, indicating that RT can serve as implicit proxy of decision confidence in our task. In what follows, the analogues of all neural effects related to confidence are shown for RT in the Supplement.

### Neural codes for choice and confidence

We trained linear decoders to predict participants’ choices or confidence judgments from the spatial activity patterns within each of 180 areas covering the cortical surface (combined across both hemispheres ^50^) at each time point (Figure 2A). Dorsal premotor cortex (PMd) and primary motor cortex (M1) exhibited the strongest choice decoding accuracy across cortex at the end of stimulus viewing (Figure 2A; PMd: 57.5% +/- 0.01%) as well as right before choice (80.4% +/-0.02%). During stimulus viewing, choice-predictive PMd and M1 activity exhibited hallmark signatures of evidence accumulation. (i) It built up with a rate that correlated with evidence strength on single trials (Figures 2B, S4A, and S5A); (ii) the cross-temporal decoding matrix^51^ was consistent with the gradual build-up of a consistent choice-coding activity pattern throughout stimulus viewing (PMd: Figure 2C, M1: Figure S4B); and (iii) the trial-to-trial variability of choice-predictive activity also grew monotonically during stimulus viewing (PMd: Figure 2D; M1: Figure S4C), a signature of diffusion processes^13,52^. After stimulus offset, the variance dropped in both regions (Figures 2D and S4C), and the cross-temporal decoding matrix showed little generalization to the stimulus interval (PMd: Figure 2C, M1: Figure S4B), both indicating a state transition, likely related to action execution. Based on these observations, we refer to the choice-predictive activity in PMd and M1 during stimulus processing as “neural DVs” in the following. Choice-predictive signals with these qualitative signatures were also observed in several areas of association cortex (Figure S5).

**Figure 2.**
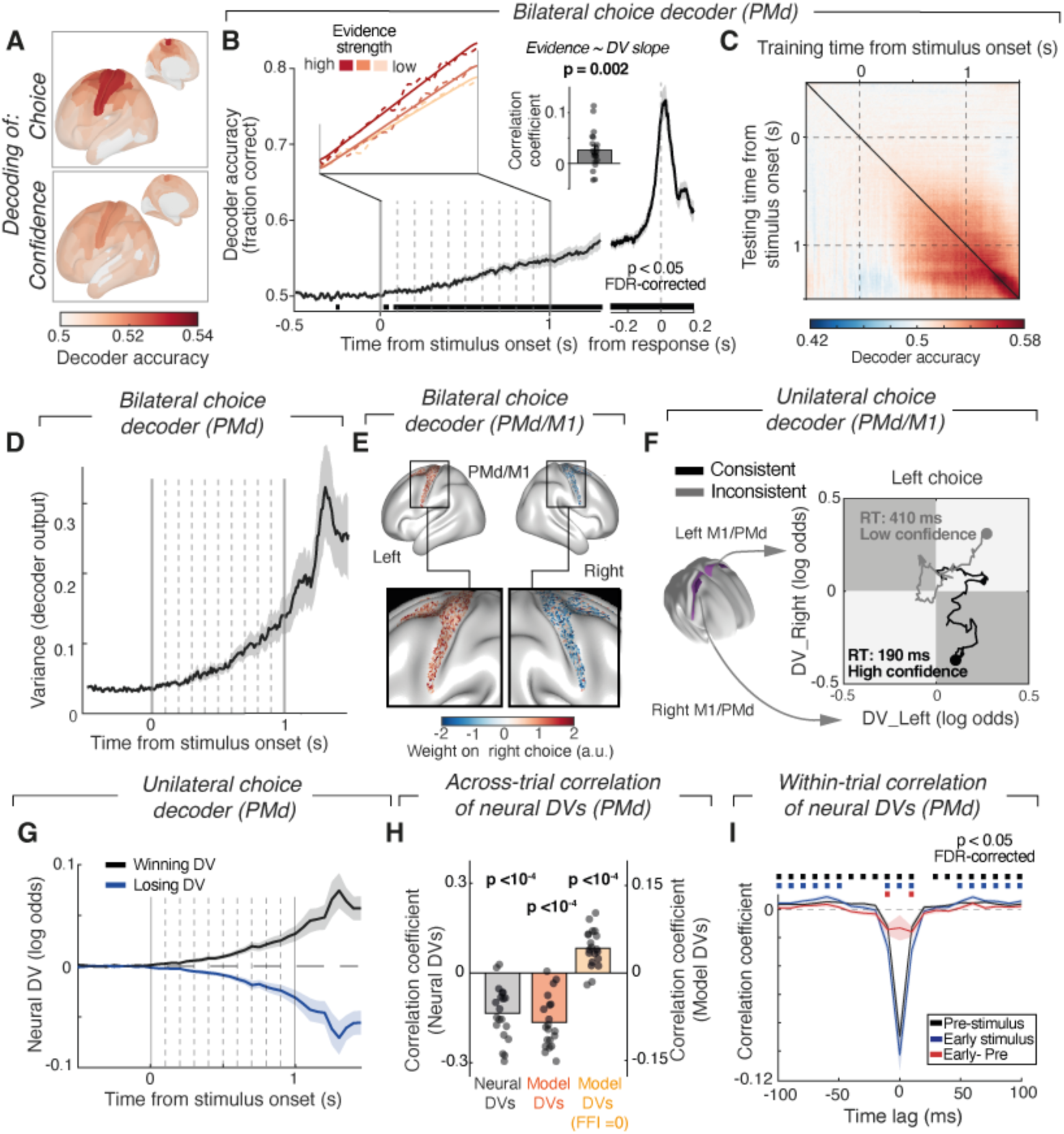
Competing neural DVs in PMd. **(A)** Whole-cortex maps of choice and confidence decoding accuracy at the end of stimulus viewing (Methods). The maps are not thresholded to highlight the full distribution of choice-selective activity. **(B)** Time-course of choice decoding from bilateral PMd, aligned to stimulus onset and choice. Top left inset: decoder accuracy time-course for three levels of evidence strength. Dashed lines, data; solid lines: fitted slopes. Top right inset: single-trial correlation between the slope of decoder output and evidence strength (Methods). Ramping slopes were fitted for the interval 0.2 to 1 s from stimulus onset (Methods). Cross-validated decoder accuracy gradually ramped up during the 1s presentation of the evidence stream, followed by a transient peak aligned to the time of the behavioral choice report. Data points, participants; p-value, two-sided permutation test. **(C)** Temporal cross-decoding matrix of choice decoder from PMd aligned to stimulus onset. Colored arrows indicate different part of the matrix reflecting different neural encoding of choice (see main text). **(D)** Variance of PMd choice decoder output. **(E)** Group average decoder weights per vertex within PMd/M1 at stimulus offset (last 100 ms). **(F)** Time courses of values of DV_left and DV_right, obtained from right and left PMd/M1, respectively, for two example trials ending with either left or right choices, for consistent trials (black lines) and inconsistent trials (gray lines). The associated RTs and confidence reports are written at the end point of the DV trajectories. “Consistent” indicates that both neural DVs point to the same choice, and conversely for “inconsistent”. **(G)** Time course of trial-averaged values of winning DV and losing DV from unilateral choice decoder applied to PMd neural activity (see main text). **(H)** Intrinsic across-trial correlation (stimulus and choice factored out) between neural DV from PMd and between the DVs of different models (with or without cross-inhibition) at the end of stimulus viewing (1 s after stimulus onset for neural DVs). DVs of model without cross-inhibition (FFI=0) follow the positive correlation of the stimulus means. **(I)** Within-trial correlation between winning and losing neural DVs from PMd time courses from two trial segments: pre-stimulus interval and early stimulus presentation (first 500 ms after stimulus onset) and their difference (in red), shown as cross-correlograms. Marks, p < 0.05 (FDR-corrected); p-values from permutation tests against 0 or between the two segments (in red). All panels: lines, group average, shaded areas, SEM. Data from N = 20 participants.

PMd, M1, and many other areas also exhibited activity predictive of confidence, with cortical distributions (Figure 2A) and dynamics (Figure S4D) resembling the choice-predictive activity. However, previous results suggest that patterns of cortical population activity encoding confidence are partially distinct from those encoding choice^53,54^. We trained classifiers to predict either choice or confidence from cortex-wide activity patterns (i.e., not separated by region). Decoding accuracy for both variables increased during stimulus viewing, with lower values for confidence (Figure S4E). The decoders for both variables (i.e., weight vectors) were nearly orthogonal early during stimulus viewing and, while becoming slightly more aligned over time, remained largely distinct toward the end (Figure S4F). In sum, the neural codes for both behavioral reports in our task were expressed in similar cortical regions, but in formats that are not linearly related. This is expected from our model in which the transformation from DVs to high versus low confidence judgments is non-linear (conditioned on the winning DV; Figure 1H). In what follows, we illuminate the transformation from the neural DVs to confidence.

### Competition between neural DVs in premotor cortex

We first focused on PMd, which previous work has implicated in decision formation for hand movement choices in monkeys^30,55^ and humans^11,27–29^. The weights of choice decoding in PMd and M1 (decoder trained on bilateral PMd/M1 activity combined; Methods) exhibited opposite polarity for both hemispheres (Figure 2E), indicating that left and right PMd and M1 both increased (decreased) their activity before contralateral (ipsilateral) choices. This lateralization is well established for M1^56^, but not for PMd, where previous studies of motor preparatory activity in monkeys reported only moderate lateralization^57^. Independently training two separate decoders in PMd/M1 of each hemisphere to predict its respective contralateral choice (Figure 2F; Methods), enabled us to simultaneously track the neural DVs for two opposing choices. In what follows, we refer to these two decoder outputs as DV_left and DV_right, respectively (Figure 2F). This two-dimensional representation provided mechanistic insight into the neural dynamics underlying decision formation within PMd/M1 (Figure 2F-I). Figure 2F shows how both neural DVs evolved over time for two example trials ending with a left choice. In one trial, labeled as “consistent”, the two neural DVs predicted the same (left) choice (black line). In the other, “inconsistent” trial, only DV_left predicted the actual left choice, whereas DV_right predicted the non-selected (right) choice option. As it happens, those two trials were associated with high (black) versus low (gray) confidence reports, and with short (black) versus long (gray) RTs – two systematic effects that we quantify below. For left choices, we label DV_left as “winning DV” and DV_right as “losing DV”, and conversely for right choices. Those two DVs exhibited the expected trial-average behavior, with the winning DV increasing and the losing DV decreasing throughout stimulus viewing (Figure 2G).

Neural circuit models of decision-making^18,24–26,37^ as well as our best-fitting accumulator model (Figure S6C) postulate inhibitory crosstalk between the two competing, choice-selective neural populations, either through feedforward and/or lateral inhibition. We reasoned, and verified with our model, that such crosstalk should generate negative correlations between the two neural populations, across and within trials. To isolate intrinsically generated DV correlations across trials, we factored out effects of stimulus category and choice by first computing separate correlations for each stimulus category and choice and then averaging the resulting correlations. The across-trial correlations between the inputs (i.e., sample means of the left and right streams) were positive (Figure S6A-B). We computed the correlations between model DVs in the same way as the neural DVs (i.e., linear decoding for left and right accumulator values, followed by correlating the decoder outputs; Methods). The DVs of the best-fitting model were negatively correlated across trials (Figure 2H, middle bar). Indeed, this was a signature of the intrinsic inhibitory crosstalk (feedforward as in our model, or feedback inhibition; see Discussion): when eliminating the cross-inhibition in the model by setting the FFI parameter to zero, the across-trial correlation between DVs turned positive (Figure 2H, right bar), now following the statistics of the stimulus input (Figure S6A-B). Critically, the neural DVs in PMd at stimulus offset were negatively correlated across trials (Figure 2H, left), indicative of cross-inhibition between the two pathways somewhere in the brain. The same was true for neural DVs in M1 (Figure S6D).

Similar results were obtained for correlations between the within-trial time courses of model or neural DVs (Figures 2I, S6E,F). Recall that the stimulus fluctuations on both sides were independent (Figure S1C, D). By contrast, the DVs in the best-fitting model were negatively correlated across a range of lags (Figure S6E), which again indicated intrinsic inhibitory crosstalk: DV correlations were absent in the model when the cross-inhibition (FFI) parameter was set to zero (Figure S5E). Note that we evaluated the within-trial cross-correlations between DVs specifically for the early stimulus intervals, because we observed that accumulator models without cross-inhibition could also show (weak) negative cross-correlations when those were computed across the whole trial, due to occasional bound crossings later in the trial (correlations were absent for models without a bound). Because bound crossings did not occur in first half of the trial, we focused on the early trial time window to obtain conservative signatures of intrinsic inhibitory crosstalk. Indeed, the neural DVs in both PMd (Figure 2I) and M1 (Figure S6F) exhibited the negative within-trial correlations predicted by our best-fitting model. Assuming that this cross-inhibition would already be in place before stimulus onset and shape the correlated variability of ongoing activity^58^, we also computed cross-correlations between neural DVs for an equally long pre-trial baseline window. Again, correlations were negative, albeit weaker than during the early trial interval (Figures 2I, S6F).

In sum, neural DVs in PMd and M1 exhibited diagnostic statistical signatures of the inhibitory interactions between “decision pathways” that are postulated by influential circuit models^25,26^ as well as our best-fitting model. Our neural data refute race models made of two independent accumulators. Our best-fitting accumulator model also well captured several other qualitative neural signatures of cortical decision dynamics including adaptation of the input encoding at the level of sensory processing in primary visual cortex (Figure S7).

### Neural DVs in premotor cortex account for confidence

The correlation between the two neural DVs in PMd implies that determining their contribution to confidence is not straightforward: if only one DV (e.g., the losing DV^19,20^) governed confidence, the other DV would inherit the link to confidence from that neural crosstalk, giving rise to a spurious association. We, therefore, capitalized on our simultaneous assessment of winning and losing DVs to relate both to single-trial confidence reports in a multiple logistic regression (Figure 3A). We also included evidence strength in the model because this variable strongly affected behavioral measures (Figure 1) as well as neural DVs (Figure 2), thus expectedly producing strong weights in the current regression (Figure 3B). Critically, we found significant weights of opposite signs for the losing and winning DVs in PMd (Figure 3C), showing that trial-to-trial variations in both competing decision signals uniquely contributed to confidence, over and above the external evidence. The absolute value of beta weights was stronger for the winning DV than for the losing DV (Figure 3D). The same effect was observed after excluding the two participants that did not show a primacy effect in the psychophysical kernels (p = 0.0002, two-sided permutation test).

**Figure 3.**
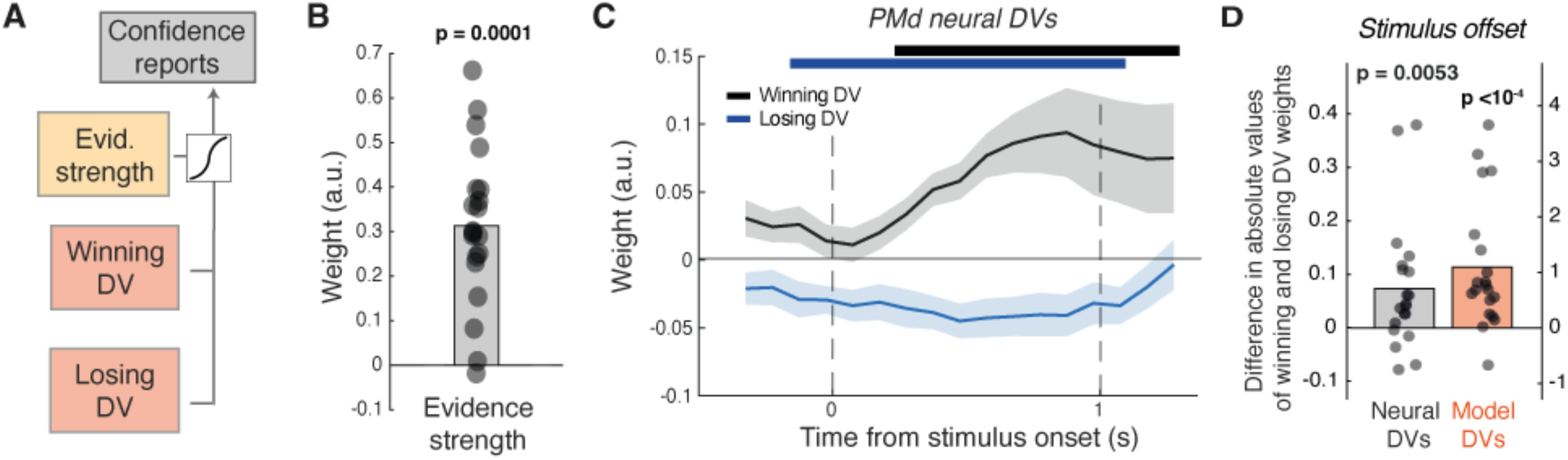
Unique contributions of competing neural DVs in PMd to confidence. **(A)** Schematic of logistic regression model predicting single-trial confidence reports from evidence strength and level of neural DVs over time (350 ms moving window, 100 ms steps, see Methods). **(B)** Regression coefficients (beta weights) for evidence strength at the end of stimulus viewing (1 s after stimulus onset). **(C)** Time course of beta weights for winning versus losing neural DVs. The latter were evaluated on a sliding-window basis (350 ms window length; Methods). Marks, p < 0.05 (cluster-based permutation test). Lines, group average; shaded areas, SEM. **(D)** Difference between absolute value of beta weights for winning and losing neural and model DVs at the end of stimulus viewing (1 s after stimulus onset). P-values in all panels are from two-sided permutation tests against 0. Bars, group average; data points, individual participants. Data from N = 20 participants.

Our best-fitting model reproduced the asymmetrical contribution of winning and losing DVs to confidence observed in PMd (Figure 3D; excluding two participants without primacy effect: p < 10^-4^, two-sided permutation test). The fitted confidence asymmetry weight was smaller than one for all participants (Figure S7I), indicating a weaker contribution of the losing DV than winning DV to confidence. Among several tested model architectures, the pattern of the neural data could only be replicated by models that included such a confidence asymmetry weight or a model, in which DVs were bounded at zero, referred to as lower reflective bound (Methods). The lower reflective bound model, which is largely consistent with realistic circuit models of decision-making^25^, fit the behavioral data worse overall (Figure S2) and captured the time course of the asymmetry in behavioral confidence kernels less well (Figure S8) than the model with confidence asymmetry weight. While it is possible that detailed circuit models will fully capture our observed behavioral and neural signatures in certain parameter regimes^25,27^, the parameters of such models cannot be fit in a principled manner.

Notably, analogous links between neural DVs in PMd and confidence reports were also found, with inverted sign, for long versus short RT trials (Figure S9A-C). The observation of correlations between winning and losing neural DVs and RT suggest that the variability of RTs in our task is governed by a process beyond evidence accumulation and bound crossing (see Discussion). An unbounded accumulation process, in which the choice governed by the larger DV, makes no prediction for RT and, therefore, for the observed correlation. In the case of an accumulation process with a fixed bound, it is possible (but not a given for our fixed duration task) that the bound-crossing time determines RT. Under this assumption, the RT variability can only be explained by variability in the level of the losing DV (Figure S9D, E, dashed line), because the winning DV reaches the same fixed bound on all trials, irrespective of the time of bound crossing. This scenario cannot explain the observed negative correlation between the winning neural DV and RT that we observed. Finally, an accumulation process with collapsing bound, combined with cross-inhibition between DVs (both of which are entailed in the best-fitting model) can predict the asymmetry in the correlations of winning versus losing DVs and bound crossing time similar to the one we observed for RTs (Figure S9D,E, solid line). However, the time courses of these model effects differ markedly from the time courses of the relationships between PMd DVs and participant RTs in our data (Figure S9C). These observations are in line with the idea that processes beyond bound crossing account for the trial-to- trial variability of RT in our task and its link to PMd activity (see Discussion).

In sum, the values of both neural DVs in PMd uniquely and asymmetrically contribute to confidence, with a stronger impact of the winning DV. This finding implies that any reduction of the two-dimensional dynamics of neural DVs into a one-dimensional signal (e.g., the difference between DVs) will lead to loss of information about confidence. Again, qualitatively identical effects to those in PMd were also found for M1 (Figure S10).

### Neural DVs in association cortex shape confidence over and above PMd

Our analyses so far focused on neural DVs with a lateralized organization, so that we could monitor competing neural DVs *within* an area. While in areas beyond PMd and M1 (and the somatomotor areas 1, 2, 3a, 3b, light gray in Figure 4A), lateralization is less strongly expressed, many of them still exhibited significant choice-predictive activity (Figure 2A, 4A). Because we here trained single decoders on activity patterns across homotopic regions from both hemispheres, we only obtained a single DV from individual areas. Excluding somatomotor cortex, a total of 22 areas (Table S1) located in frontal, parietal, and insular association cortex satisfied three criteria for neural DVs: (i) significant choice decoding at the end of stimulus viewing, over and above the choice-predictive PMd/M1 activity (Figure 5A; Methods); (ii) during stimulus viewing, a build-up of choice-predictive activity, the rate of which reflected evidence strength (Figure S5A); which is (iii) accompanied by monotonically increasing trial-to-trial variability (Figure S5B).

**Figure 4.**
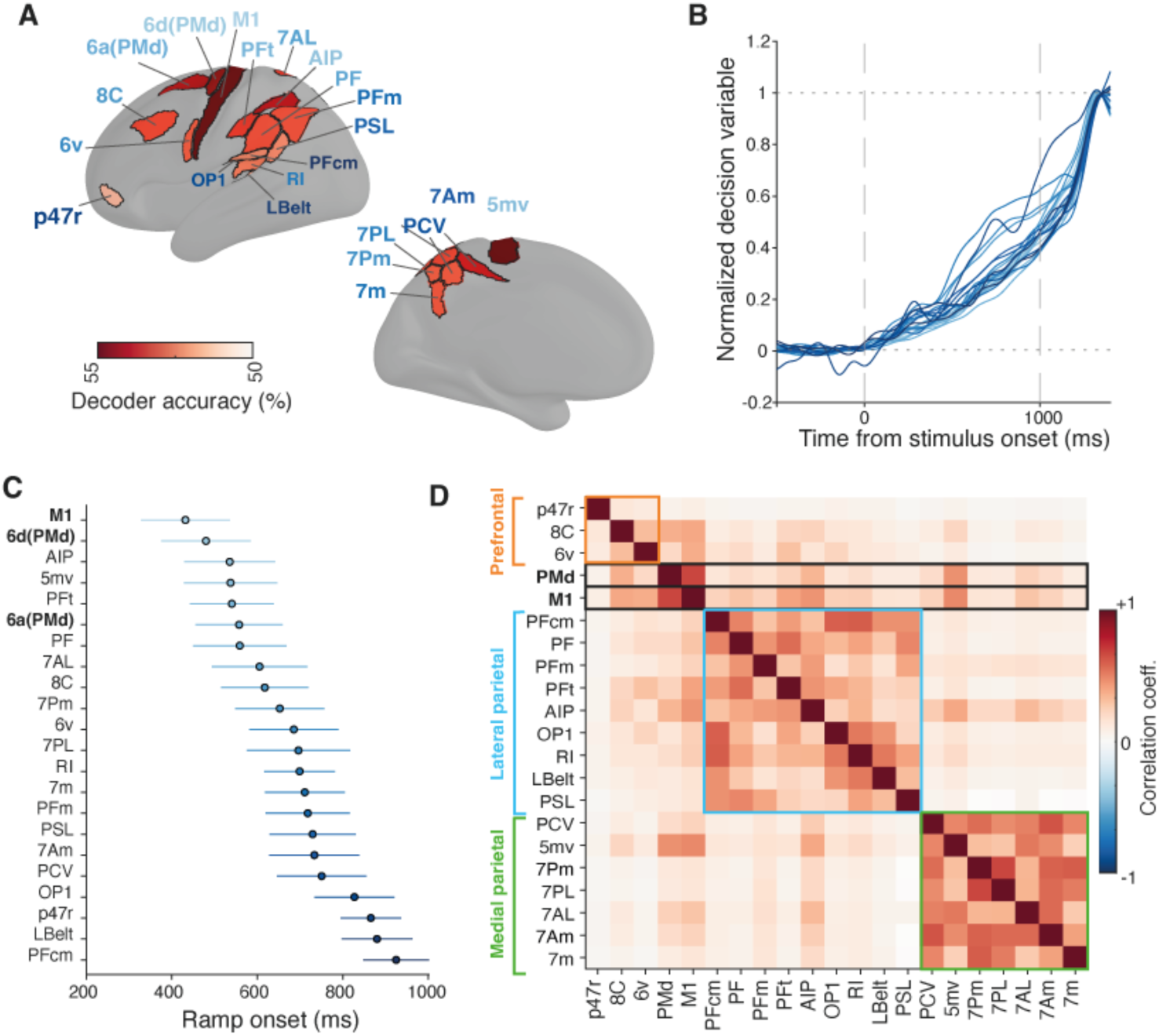
Neural DVs across association cortex. **(A)** Whole-cortex maps of choice decoding accuracy at the end of stimulus viewing for the 22 selected ROIs (excluding somatomotor cortex). (**B)** Average normalized neural DV for the selected ROIs. For each ROI, the DV was normalized by its mean value during the late decision period (1350ms), allowing comparison of temporal dynamics independently of decoding magnitude. **(C)** Estimated ramp onset for the same selected ROIs. Ramp onset was defined as the earliest time at which the local slope of the normalized DV, estimated within a 100-ms moving window, exceeded the pre-stimulus baseline by 2 standard deviations and remained positive until the peak of DV (see Methods). Points, group mean; error bars, ±SEM. **(D)** Matrix of single-trial, intrinsic correlations among the neural DVs in the final 100 ms of stimulus viewing, computed after factoring out effects of stimulus category and choice. All regions within PMd/M1 are highlighted. Data from N = 20 participants.

**Figure 5.**
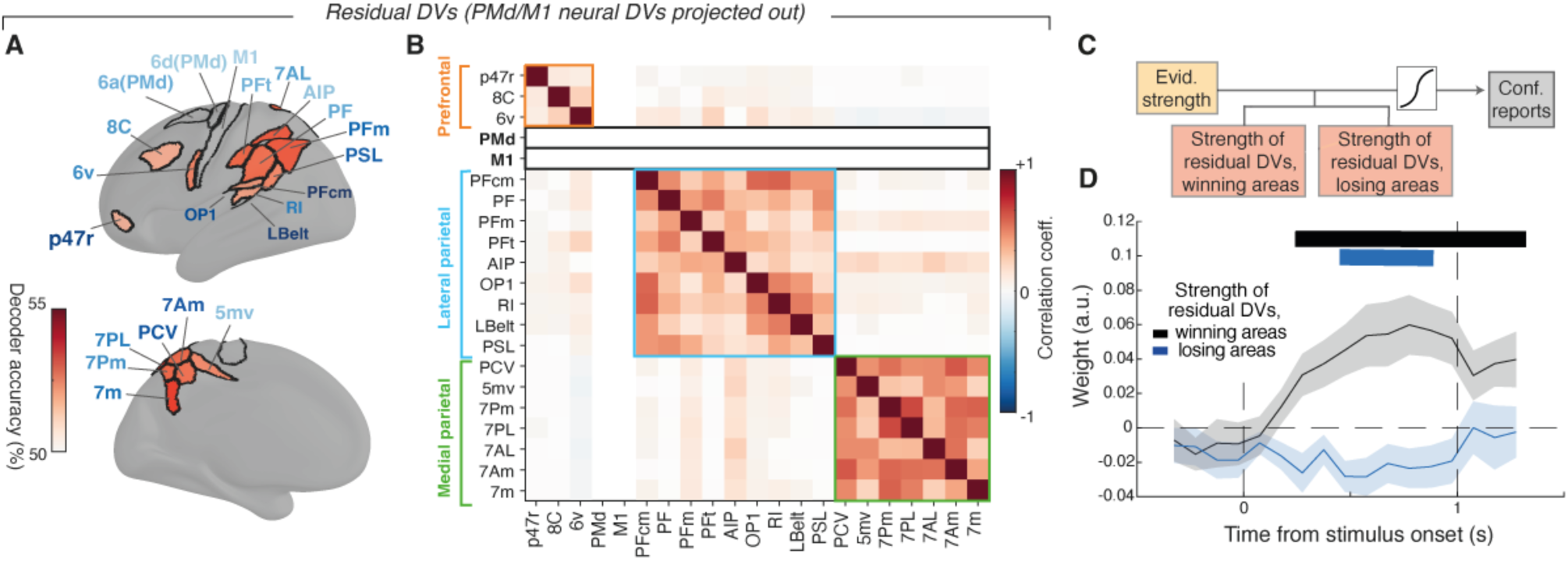
Contributions of competing neural DVs across cortical association areas to confidence. **(A, B)** As Figure 4, panels A-D, but for “residual neural DVs” after removing the PMd/M1 choice code (Methods). **(C)** Schematic of logistic regression model predicting single-trial confidence reports from evidence strength and the strength of the average residual DV from losing and winning areas after PMd/M1 DV has been projected out (see Methods). **(D)** Evolution over time of regression weights linking trial-by-trial confidence reports to the strength of the losing and winning areas residual DVs. The latter were evaluated on a sliding-window basis (350 ms window length; Methods). Marks, p < 0.05 (cluster-based permutation test). Data from N = 20 participants.

The strengths at the end of stimulus viewing of neural DVs in association cortex as well as PMd and M1 co-varied from trial to trial (Figure 4D and Figure S11A). While these neural DVs were widely distributed (Figure 4A) and intrinsically correlated (Figure 4D), they exhibited different dynamics in different areas (Figure 4B,C). Estimates of the onset latencies of neural DVs for each of the above 22 areas (Methods) suggested that neural DVs first started to ramp in the motor system (first areas: M1 and 6d within PMd, detectable from ∼432 ms after stimulus onset). Neural DVs subsequently emerged in posterior parietal area AIP (mean latency: 535 ms) and prefrontal region 8C (mean latency: 617 ms; Figure 4C). In other words, neural DVs were strongest and earliest in human motor and dorsal premotor cortex in our task.

Regardless of the origin of these DVs in association cortex, whether they were computed locally or inherited from PMd/M1, we reasoned that DVs in association cortex may contribute to confidence over and above the PMd/M1 DVs. Given the correlation of PMd/M1 DVs with confidence and with the DVs in association cortex (Figure 4D), the relationship between the association cortical DVs with confidence may be spurious, due to genuine coupling between neural DVs and/or spatial leakage of the activity estimates of PMd/M1 and any of the association regions. To remove such spurious correlations, we removed the component that each of the 22 association cortical DVs shared with the PMd/M1 DV (Methods and Figure S11A-B). The resulting “residual” DVs were, while less predictive of choice (Figure 5A), still statistically significant and remained coupled across many region pairs, with a structure highlighting three clusters: in prefrontal, lateral parietal, and medial parietal cortex (Figure 5B).

We classified the residual DVs of all regions as “winning” or “losing”, depending on whether they supported the selected or non-selected option on that trial. This was done for each of many windows sliding across the trial. We then computed the strength of the support for the selected option as the mean of the absolute values of all winning DVs and, likewise, the strength of support for the non-selected option as the mean of the absolute values of all losing DVs. Those measures, whose correlation was negligible (Figure S11D), were then regressed on single-trial confidence reports (logistic) or RTs (linear). Again, the regression models contained evidence strength as co-regressor, to ensure that effects between neural DVs and confidence were not accounted for by the effect of evidence strength (Figure 5C; Figure S11E).

We found a positive relationship between confidence and the strength of winning DVs across association cortex and a negative relationship with the strength of losing DVs (Figure 5D). This effect built up during stimulus viewing, peaking at around 750 ms from stimulus onset. The same effect, with the expected flipped sign, was observed for RT (Figure S11F). These results indicate that confidence reflected how widely distributed decision signals beyond PMd/M1 activity favored one option over the other.

## Discussion

The current work was motivated by the growing realization that neural populations selective for distinct choices are distributed across many regions of the cerebral cortex^12,33–36,59^. Building on theoretical insights^21,60^, we reasoned that such distributed cortical decision dynamics may shape the sense of confidence associated with a behavioral choice. Due to technical limitations, it has been so far challenging to directly and simultaneously monitor such cortex-wide neural competition during decision formation. We developed an approach to overcome that challenge and identify statistically unique links between decision-related neural dynamics within several cortical areas and confidence. Our approach provided several new insights. First, neural DVs encoding opposite choices in PMd are intrinsically negatively correlated, which modeling identified as a signature of neural cross-inhibition in the context of our task. Second, neural DVs in the motor system (areas M1 and PMd) were strongest and emerged early during deliberation. Third, both winning and losing DVs within PMd made unique but asymmetric contributions to confidence, with a dominance of the winning DV. Fourth, trial-to-trial variations in neural DVs in parietal and prefrontal association cortex over and above the DVs measured in PMd/M1 predicted choice and explained part of the trial-by-trial variation in confidence. Together, our results are consistent with the idea that confidence is shaped by the ongoing competition during deliberation between many neural DVs distributed across the cortex.

Theoretical work has proposed that perceptual evidence accumulation is implemented through recurrent interactions between multiple areas, rather than a single processing stage within a feed-forward pathway^34,38^. While some work in mice has shown that several of these distributed regions are necessary for evidence accumulation and decision-making^61^, other findings from rats indicate that decision signals first emerge in a small set of frontal and striatal regions implicated in action planning, followed by decision signals in association cortex^35^. Some of our current human results resemble the latter, with the cortical motor system leading the build-up of decision signals. Regardless of whether evidence is accumulated in a distributed network, or first within (pre-)motor cortex and subsequently fed back to association cortex, our results indicate that the brain uses the decision variables carried by multiple, spatially distinct neural populations within PMd/M1, prefrontal, and parietal association cortex, and winning as well as losing populations in these areas, for computing confidence. Establishing a causal role of these distributed choice codes for confidence formation will require a combination of spatially specific perturbations with MEG recordings that is currently not affordable.

Neural signals other than those linearly related to such choice codes, possibly expressed in other brain areas, may also contribute to the sense of confidence^53,62,63^. In line with this idea, we found that our large-scale (cortex-wide) decoders for confidence and choice, while showing substantial overlap in the cortical distributions of decoder accuracy, used activity patterns (i.e. weight maps) that were only weakly aligned. One possibility explaining this local misalignment between neural choice and confidence codes is that confidence codes are computed from the evidence through a distinct process from the one that computes choice codes. Another possibility is that local choice codes are transformed into local confidence codes in a non-linear manner (e.g., rectification or the conditional transformation postulated by our model) and/or by pooling choice signals from regions remote from the one carrying the confidence code. This scenario is in line with our observation of multiple local “winning” and “losing” neural DVs contributing to the confidence reports. A final possibility, which is not mutually exclusive with the previous ones, is that many confidence signals detectable in cortical population activity do not reflect the neural code that drives the behavioral confidence report, but are secondary signals, for example reflecting neuromodulatory signals broadcasting overall confidence or uncertainty^1,4,64,65^.

Influential algorithmic models of decision-making have assumed a key contribution from the losing accumulator to confidence judgments^19,20,66,67^. In these models, the winning decision variable typically, has no contribution because it is clamped at the constant value of the decision bound on all trials, so that trial-by-trial variations of confidence can only result from variations of the losing decision variable^19,66,67^ (or its combination with elapsed time ^20^). By tracking these two neural races simultaneously in human frontal cortex, we here show that both DVs make unique contribution of opposite polarity to confidence (over and above evidence strength), with a stronger impact by the winning neural DV. Our model explains why the impact of the winning DV was not zero: a collapsing bound induces time-dependent trial-by-trial variation of the level of the winning DV at bound crossing. This, combined with a weight applied to the losing DV was sufficient in our model to match the distinct impact of each DV.

Our ability to simultaneously track the two competing neural DVs depended on the lateralized format of choice-selective preparatory activity in human PMd and M1. The maps of the weights of our linear decoders clearly support this lateralization of action-selective motor preparatory PMd activity in our data. We also found the same pattern of confidence associations for both areas. Even so, the decoders trained separately on PMd and M1 activity in both hemispheres are likely to each contain a mixture of neurons selective for contralateral and ipsilateral hand choices, with a dominance of contralateral-preferring neurons. Furthermore, these two neural groups with preference for contra- and ipsilateral choices are likely to process a mixture of the inputs from both contra- and ipsilateral evidence streams. This mixture results from the cross-inhibition between the processing pathways for both sides postulated by our winning models, as well as from only partial hemispheric lateralization of the sensory input processing of the competing neural races. Here, we conceptualized the two decoder outputs as population-level proxies of the state of two competing neural accumulators.

We interpret the negative correlation between left and right PMd decoders as a signature of intracortical inhibitory interactions. In behavioral tasks often used for studying evidence accumulation^10,27,42,43^, evidence for the choice alternatives is (partially or perfectly) correlated: a sample supporting one choice provides, at the same time, evidence against the alternative^44^. Thus, any negative correlations at the level of neural DVs may be induced by the input and cannot be interpreted as evidence for intrinsic competition. Our task instead eliminated such extrinsic correlation. While the primary motivation for this design choice was to enable the identification of competition intrinsic to the brain, we note that largely independent noise fluctuation of the evidence supporting different options may be a common feature of many real-world decision problems, including foraging for rewards^68^. The negative crosstalk between decision variables took the form of feed-forward inhibition in the model that best fit the behavior of our participants. There is evidence for both feedforward inhibition (in sensory cortex^69^) and for feedback inhibition (in action-related, frontal regions^70–73^) in the cerebral cortex. We speculate that both forms of inhibition conspired to shape cortical dynamics and behavior in our task.

Our results partly converge with findings from studies linking neural population dynamics in monkey lateral intraparietal (LIP) area^54,74^ or dorsolateral prefrontal cortex^75^ to implicit or explicit measures of confidence. These studies decoded the probability of the decision being correct^74^ or the decision confidence reported by monkeys^54^ from LIP activity, with a time course similar to that of the time course of the decoding of the concomitant choice. In contrast to our work, these results indicated that the losing population makes only a minor contribution to confidence^54,74^, with heterogeneity within the winning population explaining confidence variations^74^. This difference to our conclusions may reflect a genuine difference between species and/or the nature of confidence computations in parietal versus frontal cortex. However, neither of these previous monkey studies assessed the correlation between winning and losing neural DVs to determine their statistically unique contributions to confidence.

Bounded accumulation is analogous to models that explain confidence judgments by the combination of DV and elapsed time^16,20^ and approximates a Bayesian process that updates its belief about the underlying category upon new samples and commits to a choice when the value of expected accuracy increase is less than the cost of processing further evidence^17^. In our best-fitting model, the source of the confidence asymmetry occurs at the readout stage of processing, and truncation of DVs at a lower bound was not sufficient to capture the observed patterns. This model replicated key signatures of the neural decision variables in PMd. However, the above-mentioned Bayesian belief updating computation (partially-observable Markov decision process) can produce the confidence asymmetry observed in the psychophysical kernels without any re-weighting of evidence at the readout stage^17^. Future work should elucidate how such a process can be approximated in neural dynamics made up of at least two competing and weakly coupled neural populations, and whether this can account for our findings.

A limitation of our fixed duration task is that the time of decision commitment was not well identified. Our best-fitting accumulator model contained a collapsing bound, which is in line with a normative model (partially observable Markov decision process) for fixed duration tasks that postulates a cost of gathering further evidence^17^. We estimated the time of bound crossing from many simulations of the model dynamics for each trial under different realizations of noise, which reproduced several qualitative patterns observed for participants’ RTs. However, our behavioral data suggest largely separate sources of variability underlying RTs and confidence reports, which were only weakly correlated across the group, and not at all for some individuals. While recent rodent work established model-based analyses of cellular-resolution neural data to infer the time of decision commitment in fixed duration tasks^35,76^ such approaches will need to be carefully validated before being applicable to non-invasively recorded neural mass signals that are noisier. Either way, the observation that participants’ RTs and neural signals related to RT mirrored all qualitative signatures of confidence reports is noteworthy. The additional RT variability underlying these signatures may originate from action-related circuits, potentially also involving basal ganglia. Indeed, measures of response vigor also exhibit signatures of decision confidence both in humans and rodents^77–79^. The RT correlates of confidence observed here could reflect the impact of the DVs at the commitment time on response vigor. It will be important to determine the sources of additional RT variability by combining our task with measurements of sub-threshold muscle activity and/or using a free response version of our task.

Neuroscience has recently started to address the distributed nature of cognitive computation^60^. Our current study contributes to this nascent field by identifying several signatures of distributed neural computation related to confidence. Future successful circuit models of the brain-wide neural dynamics of decision and confidence formation will need to reproduce these neurophysiological signatures.

## Methods

### Participants

Twenty-three healthy human participants (mean age 28, range 21-40, 9 females) took part in the study after informed consent and introduction to the experimental procedure. The study was approved by the ethics review board of the Hamburg Medical Association responsible for the University Medical Center Hamburg-Eppendorf (UKE). Exclusion criteria, all of which were assessed by self-report, were: history of any neurological or psychiatric disorders, hearing disorder, history of any liver or kidney disease or metabolic impairment, history of any chronic respiratory disease (e.g., asthma), history of hyperthyroidism or hypothyroidism, pheochromocytoma (present or in history), allergy to medication, known hypersensitivity to memantine or lorazepam, family history of epilepsy (first or second degree relatives), family history of psychiatric disorders (first or second degree relatives), established or potential pregnancy, claustrophobia, implanted medical devices (e.g., pacemaker, insulin pump, aneurysm clip, electrical stimulator for nerves or brain, intra-cardiac lines), any non-removable metal accessories on or inside the body, having impaired temperature sensation and / or increased sensitivity to heat Implants, foreign objects and metal in and around the body that are not MRI compatible, refusal to receive information about accidental findings in structural MR images, hemophilia, frequent and severe headaches, dizziness, or fainting, regularly taking medication or have taken medication within the past 2 months. Three participants were excluded from the analyses: one due to excessive artifacts in the raw MEG data resembling metal artifacts, and the other two due to not completing all six experimental sessions. Thus, we analyzed and report the results from 20 participants (7 females, 13 males), except where explicitly noted (several results are also reported after excluding the 2 participants who did not show primacy in the psychophysical kernels; Figure S1F). Participants were remunerated with 15 Euros for the behavioral training session, 100 Euros for each MEG session, 150 Euros for completing all six sessions, and a variable bonus, the amount of which depended on task performance across all sessions. The maximum bonus was 150 Euros.

### Experimental design

The study was part of a larger study that spanned seven experimental sessions and also included other behavioral task conditions as well as low-dose pharmacological interventions, all of which are beyond the scope of the present paper. The following substances were administered orally, in two sessions each (order counter-balanced; about 150 min before start of MEG recordings), in a double-blind, randomized, crossover design: the NMDA-receptor antagonist memantine (15 mg; clinical steady-state daily dose is 20 mg), the GABA-A receptor agonist lorazepam (1 mg; common clinical daily dose ranges from 0.5 to 2 mg), or a mannitol-aerosil placebo, all encapsulated identically to render them visually indistinguishable. Each session also entailed a 10 min resting-state block and a simple auditory task. The pharmacological effects on these tasks have been described in previous work^80,81^. As expected from the low dosages, choice behavior in the current task was qualitatively similar across pharmacological conditions, with subtle effects on contrast threshold (slightly increased under both drugs) and RTs (slightly prolonged under lorazepam), but no statistically significant effect on the fraction of high-confidence reports (Figure S12). Because of these subtle effects on overall behavior, and the current focus on the asymmetric contributions of competing neural DVs to confidence reports, the analyses presented in the current paper aggregated across conditions. The six sessions were scheduled at least one week apart to allow plasma levels to return to baseline (plasma half-life of memantine: ∼60 to 70 h ^82^; half-life of lorazepam: ∼13 h^83^). Each MEG session was around 2.5 hours in duration and consisted of a resting state-block and two behavioral tasks including the one described below that is the focus of the current study.

### Stimulus and behavioral task

Each trial of the task consisted of the following sequence of events (Figure 1A). After a variable delay (uniform between 0.5 and 1 s) ten successive samples of variable contrasts (100 ms each) were shown in each visual hemifield. Participants were asked to compare the average mean contrast across the left and right samples (i.e., forced choice report: “left is stronger” or “right is stronger”). The offset of the last sample marked the beginning of the response period for participants. Participants reported their binary choice (left vs. right), and their confidence about the correctness of that choice simultaneously, by pressing one of four different buttons, whereby the two hands were always mapped to different choices. The index and ring fingers of each hand were then used to report confidence. During MEG sessions, participants used two response pads, one for each hand. During the training sessions participants, used the same stimulus-response mapping, but pressed keys on a computer keyboard. After a participant’s response and a consecutive variable delay between 0 and 0.5 s auditory feedback was given (250 ms duration). A low tone indicated a wrong answer, and a high tone indicated a correct answer. The ten consecutive contrast samples from the two hemifields were drawn from two normal distributions of contrast levels whose mean difference (right - left) was centered on each participant’s 75% accuracy contrast level. This threshold was determined by running a QUEST staircase continuously in the background. The standard deviation of each of the two normal distribution was 0.15. After each block of 120 trials participants could take a short self-timed break. After the third block, participants took a longer break lasting at least five minutes. Each participant completed seven sessions, whereby each session consisted out of six blocks of 120 trials, lasting approximately 90 min. The first session was a training session that took place in a behavioral laboratory and was used to expose participants to the task and to calibrate their performance to 75% correct. The subsequent six sessions were experimental recording sessions that took place in the MEG laboratory and yielded the data analyzed in this paper.

### Data acquisition

#### Behavior and eye tracking

Stimuli were generated using Psychtoolbox-3 for MATLAB and were back-projected on a transparent screen using a Sanyo PLC-XP51 projector at 120 Hz during MEG recordings, or on a VIEW-Pixx monitor during the training session in a behavioral psychophysics laboratory. Eye movements and pupil diameter were recorded at 1000 Hz with an EyeLink 1000 Long Range Mount system (equipment and software, SR Research).

#### MEG

We used a CTF MEG system with 275 axial gradiometer sensors and recorded at 1200 Hz, with a (hardware) anti-aliasing low-pass filter (cutoff: 300 Hz). Recordings took place in a dimly lit magnetically shielded room. We concurrently collected eye-position data with a SR-Research EyeLink 1000 eye-tracker (1000 Hz). We continuously monitored head position by using three fiducial coils. After seating the participant in the MEG chair, we created and stored a template head position. At the beginning of each following session and after each block we guided participants back into this template position. We used Ag/AgCl electrodes to measure ECG and vertical and horizontal EOG. During MEG recordings, participants were seated on a chair inside the magnetically shielded room.

#### Magnetic resonance imaging

Structural MRIs were obtained for each participant. We collected T1-weighted magnetization prepared gradient-echo images (TR = 2300 ms, TE = 2.98 ms, FoV = 256 mm, 1 mm slice thickness, TI = 1100 ms, 9° flip angle) with 1 × 1 × 1 mm^3^ voxel resolution on a 3 T Siemens Magnetom Trio MRI scanner (Siemens Medical Systems, Erlangen, Germany). Fiducials (nasion, left and right intra-aural point) were marked on the MRI.

### Behavioral data analysis and modeling

Behavioral data were analyzed, and computational models implemented and fitted, using customized MATLAB code.

#### Psychometric curves

We modeled the psychometric function depicted in Figure S1C as:

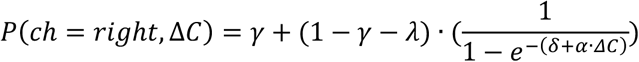

Where *γ* and *λ* are the probabilities of stimulus-independent errors (i.e., lapses), *ΔC* is the delta contrast (right-left), *α* is contrast sensitivity and *δ* is a bias term. The parameters were estimated by minimizing the negative log-likelihood using MATLAB *fmincon* function.

#### Psychophysical kernels

Psychophysical kernels were computed by comparing the two distributions of single-trial sample delta contrast (right-left) values obtained for each position of the sequence, when conditioning on each behavioral choice. This was done separately for each stimulus category (right stronger or left stronger). Specifically, we computed the receiver-operator-characteristic curve (ROC), separately for each sample position, and computed the area under the ROC curve as a measure of separability between the two distributions. The resulting two AUC time courses, one per stimulus category, were then averaged, yielding a single psychophysical kernel (Figure S1E).

#### Evaluation of confidence dependence on contrast fluctuations

We quantified the impact of contrast fluctuations on confidence by computing the average excess contrast over the mean of each stimulus generative distribution, separately for the selected and non-selected side (winning vs losing) and for each confidence report. In Figure S3E-F we quantified the impact of contrast fluctuations on RT by computing the average excess contrast over the mean of each stimulus generative distribution, separately for the selected and non-selected side (winning vs losing) and for short and long RT trials, defined as trials in which RT was shorter or longer than the median RT obtained in each session.

#### General model architecture

The general model architecture consisted of three modules transforming the two streams of contrast samples into choice, confidence reports, and associated RT.

##### Sensory module

The contrast input values *I_L_* and *I_R_* were multiplied with a sensory adaptation kernel to provide the instantaneous evidence *E_L,n_* and *E_R,n_* . The adaptation kernel consisted of an exponentially decaying function with an offset parameter (normalized as to sum to 1):

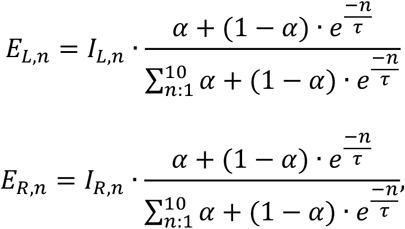

where n was sample number, and time constant *τ* (0-15) and offset *α* (0-1) were free parameters.

##### Accumulator module

The core of the model was formed by two leaky accumulators modeled by the two decision variables DV_L_ and DV_R_ with a free leak parameter, and which competed through feedforward inhibition. Accumulator dynamics were governed by the following equations:

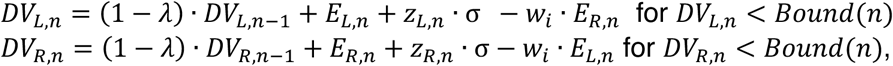

where *λ* (range -1 to 1) was a leak parameter which can be positive implying an effective leak (i.e. decay towards zero of the DVs) or negative implying an intrinsic growth of the DV towards the boundary. The noise variables *z* _L,*n*_ and *z* _R,*n*_ were random independent variables distributed as *N*(0,1) and the parameter *σ* set the magnitude of this Gaussian noise (0 to 0.05). The weight *w_i_* set the magnitude of the feed-forward inhibition (range: 0 to 1). A bias parameter B (range from -10 to 10) was added at each *DV_L_*_,*n*_ and subtracted at each *DV_R_*_,*n*_.

##### Absorbing decision bound

The model contained a collapsing decision bound of the following form: 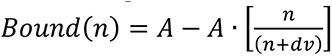,

where A was the bound starting point (range: 0 to 15), and dv was its rate of decay (range: 0 to 50). The decision time of the model was estimated as the mean of bound crossings time from 1000 simulations of each individual trial.

##### Response readout module

The model choices were governed by the accumulator that first reached the decision bound or by the accumulator with the larger value at stimulus offset. A model confidence value was computed as the difference between the winning and the losing model DV whereby the losing accumulator output was weighted by a free parameter (confidence asymmetry weight, range 0 to 1). Note that the transformation from model DVs to the confidence value was non-linear (akin to rectification) because the subtraction was conditioned on the winning and losing side, so that confidence values could only be positive. The model confidence value *Conf_model_* was transformed into the probability of a high confidence report via the logistic function: 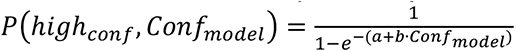 with free parameters a (range: 0 to 15) and b (range: 0 to 10).

#### Model variants

We tested a total of nineteen models of the general form described below, but differing in complexity and parameterization. Models 1 to 3 did not entail multiplication with any sensory adaptation kernel, effectively removing any transformation of the external input. In Models 1, 4 and 7, *w_i_* was set to 0, removing inhibition from the dynamics. Models 1,4,7,12,15 did not contain any bound.

In Models 2, 5, 8, 13, 16 the accumulation terminated when one accumulator reached a fixed bound A (range: 0 to 15). The lateral inhibition contained in Models 10-11 and 15-17 and 19 was as follows:

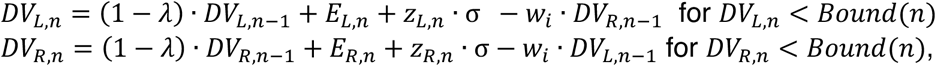

where *w_i_* (range 0 to 1) now governed the strength of inhibition from one to the other accumulator. In the brain, neural responses are bounded from below. Therefore, in Models 12-17, we also applied rectification to the DVs, so that any negative values were set to zero. In Model 18 and 19, we instead fit a reflecting lower bound as a free parameter.

#### Model fits and model comparisons

For each model the optimal parameters were found using Bayesian Adaptive Direct Search (BADS) method ^84^. At each trial, the probability for each choice and confidence report combination to be selected by the model was calculated (*P_Ri_*_gℎ*tc*ℎ,*Hi*gℎ*conf*_; *P_Leftc_*_ℎ,*Hi*gℎ*conf*;_; *P_Ri_*_gℎ*tc*ℎ,*Lo*w*conf*;_; *P_Leftc_*_ℎ,*Lo*w*conf*;_) across 1000 simulations. The log-likelihood that was maximized by BADS was calculated as:

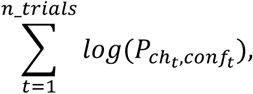

where *P_c_*_ℎ*t*,*conft*_ was the probability for the model to give as an output the same choice and confidence as the one in trial *t*. Each fitting session started by generating 500 random parameter sets, drawn from a uniform distribution bounded by the range of each parameter. We then computed the likelihood of the model parameters given the data. The parameter set with the best fit was used as the starting point for optimization routine. We used a 5-fold cross-validation procedure. Trials were divided into five equally populated subsets, separately for high and low confidence trials, so that each subset kept the participant’s proportion of high and low-confidence reports. In each fold, four subsets were used for fitting, and the remaining subset was held out for testing. The parameters fitted in each fold were then used to compute the log-likelihood of the held-out subset.

To compare the goodness of fit of the nineteen tested models, we computed for each subject (i) the average negative log likelihood across held-out trials, (ii) the mean squared error between model simulations and data of the average excess contrast over the generative mean, calculated separately for the selected and non-selected side (left or right) and high and low confidence report, as depicted in Figure 1G (iii) the Exceedance probability computed as in reference^85^.

#### Model recovery

To assess the recoverability of our best-fitting model, we generated 150 synthetic datasets from that model and fitted all candidate models considered to these data. The inputs were identical to the contrast values for each participant in our experiment: on each trial, the samples on the stronger and weaker side were drawn from Gaussian distributions (m = 0.5 ± *ΔC*/2; s = 0.15), whereby the contrast differences *ΔC* matched the values generated by the staircase values seen by the participant. Model parameters were drawn uniformly from the ranges used to initialize the real fits. We then fitted each of seven candidate models to every synthetic dataset. For each dataset, we selected the model with the lowest Bayes Information Criterion (BIC) as the winning model.

#### Model prediction of RTs and confidence reports

We quantified how well participants’ RTs and confidence reports could be predicted from the following model architectures described above: model 8 (fixed bound), model 9 (best fitting model), and model 14 (with rectification). For all models the best parameter set of each subject from the behavioral fits was used. Differently from the model fitting, the time resolution of the accumulation process was increased by ten times. We used regression models to test how trial-by-trial variations in model DV related single-trial RTs or confidence reports. The design matrices of the models contained evidence strength (absolute value of delta contrast) and the winning and losing model DV as three regressors.

### MEG data analysis

MEG data were analyzed with a combination of custom-written MATLAB code and the following toolboxes: FieldTrip^86^ version 20201009 for MATLAB (2021a; The Math-Works, Inc., Natick, MA) as well as MNE^87^ and pymeg for Python (https://github.com/DonnerLab/pymeg) established in previous work from our laboratory^11^.

#### Preprocessing

We used an automated algorithm to label artifacts in the continuous time series recorded in each session. Sensor jumps were detected by convolving each sensor with a filter designed to detect large sudden jumps and subsequently by looking for outliers in the filter response. Muscle and environmental artefacts (e.g., cars passing by the vicinity of the MEG room) were detected by filtering each channel in the 100–140 Hz or <1 Hz range, and by detecting outliers that occurred simultaneously in many channels. After removing epochs containing head movements, squid jumps and muscle artifacts, the remaining time-series were then subjected to temporal independent component analysis (infomax algorithm), and components containing blink or heartbeat artifacts were identified manually and removed. We applied a notch filter to remove 50 Hz power line noise and its harmonic (100-150 Hz) and a high-pass linear phase FIR filter (order 3) with cut-off frequency of 0.1 Hz. The resulting data were segmented in trial epochs of 2 s duration (0.5 s baseline) time-locked to stimulus onset and all epochs that contained artifacts as defined above were discarded. Finally, the preprocessed data were down sampled to a sampling rate of 400 Hz.

#### Source reconstruction

We used linearly constrained minimum variance (LCMV) beamforming^88^ to project the MEG data into source space. To this end, we constructed individual three-layer head models (inner skull, outer skull, and skin) from each individual subject’s structural MRI scans. These head models were aligned to the MEG data by a transformation matrix that aligned the average fiducial coil position in the MEG data and the corresponding locations in each head model. Transformation matrices were then generated. We computed one transformation matrix per recording session. We then reconstructed individual cortical surfaces from the structural MRIs and aligned the Glasser atlas^50^ to each surface. Based on the head model, we generated a forward model (“leadfields”) for the computation of LCMV filters that was confined to the cortical sheet (4096 vertices per hemisphere, recursively subdivided octahedron). To compute the LCMV filter for each vertex, the leadfield matrix for that vertex was combined with the trial-averaged covariance matrix of the (cleaned and epoched) data estimated for the stimulus interval (0 to 1 s from stimulus onset). We chose the source orientation with maximum output source power at each cortical location. We used an established anatomical parcellation of the human cortical surface^50^ to define regions of interest (ROIs) for further analyses, which included left and right primary visual cortex for the assessment of stimulus codes and a number of downstream areas for the assessment of neural DVs (see section “Selection of ROIs for analysis of neural DVs” below). We combined dorsal premotor cortex (areas 6a and 6d) and primary motor cortex (area 4) into the composite ROIs labeled as “PMd/M1” throughout this paper, in keeping with monkey physiology work on perceptual decisions reported via hand movements^30^.

#### Decoding of stimulus, choice, and confidence

##### Extraction of decoding features

We extracted broadband activity from each individual vertex within the selected ROI, for each time-point t and subject. The resulting vertex time series were baseline-corrected by subtracting the single-trial pre-stimulus activity (mean of interval: -0.5 - 0 seconds from stimulus onset) and then z-scored across trials. Dimensionality reduction was performed using Principal Components Analysis (PCA). We kept all PCs that cumulatively explained 90% of the variance.

##### Stimulus decoding

We used cross-validated linear regression to decode individual contrast samples from source-reconstructed broadband activity within the part of V1 located in the hemisphere contralateral to the respective stimulus sample. We used the projection of the activity on the selected PCs at each time-point t to predict the contrast of each sample i (i ɛ{1..10}). For each subject, region of interest and contrast sample position, we fitted the regression model on each time point. The projections on the selected PCs were z-scored based on training data only and the same transformation was applied to test data before evaluating prediction performance. We evaluated prediction performance within each participant, by computing the Pearson correlation across trials between predicted and presented contrast at each time point. We finally averaged the resulting time courses of correlation coefficients across participants.

##### Choice and confidence decoding from ROI activity

We used cross-validated logistic regression classifier to decode binary choice and confidence reports from source-reconstructed broadband activity within different ROIs. The choice decoder was applied to activity patterns combined across homotopic ROIs from the left and right hemispheres, or, for the analysis of DV_left and DV_right in PMd/M1, separately to left or right hemispheric ROIs. First, we extracted broadband activity from each individual vertex within the selected ROIs, for each time-point and subject. The signal was baseline-corrected by subtracting the single-trial pre-stimulus activity. Due to large number of features, we performed dimensionality reduction after z-scoring of neural activity. We then computed principal component analysis (PCA) across trials, separately for each time-point *t*, and then selected all principal components (PCs) that cumulatively explained 90% of the variance. These PCs were used as features for the decoder to predict choice. We fitted the model separately for each subject and each region of interest, at each time point. We used tenfold cross-validation within each session: trials were randomly split into ten equal subsets (folds), and in each iteration the decoder was trained on nine folds and tested on the held-out fold. Selected PCs were z-scored based on training data only and the same transformation was applied to test data before evaluating prediction performance. Since choices and confidence reports were not equally distributed, we up-sampled the minority class in the training set by randomly repeating elements until the frequency of choices or confidence reports was equal. We evaluated prediction performance by computing the percentage of correct predictions. Neural DVs were defined by extracting the single-trial log-odds obtained from the logistic regression binary classifier. In all decoding analysis, the decoder was trained and tested on the same time-point, while for computing across-time generalization matrices, all combinations of train and test time-points were used.

##### Large-scale decoding of choice and confidence from cortex-wide activity

We trained decoders to predict choice or confidence from the activity pattern across all cortical vertices that were parcellated into different ROIs for our main analyses. We extracted the broadband activity from each individual vertex within all ROIs, for each time-point and participant. The broadband signal was baseline-corrected by subtracting the single-trial pre-stimulus activity. We then computed PCA across trials, separately for each time-point and then selected all PCs that cumulatively explained 90% of the variance. We then trained logistic regression models to predict choice or confidence from the resulting PCs (number ranging from 11 to 49), using the same cross-validation procedure described above for the ROI-specific decoding. For each fold and time point, we computed the cosine similarity between the weight vectors of the two decoders as follows:

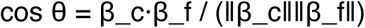

#### Spatial weight maps for unilateral PMd/M1 choice decoders

For computing spatial weight maps, the above, cross-validated logistic regression on choice was fitted using the (baselined and z-scored) single-trial activity time courses from each individual vertex within PMd/M1 without any dimensionality reduction. The beta coefficient obtained for each vertex were then extracted for each subject.

#### Transformation of model DVs into log-odds units

To generate model predictions for the observed correlations between neural DVs (see below), the model DVs were transformed into the same units (log-odds) as the neural DVs. We trained the same logistic regression models described above for MEG choice decoding to predict single-trial model choices time point by time point from the simulated model DVs. We trained the logistic model separately to each accumulator DV, to predict the choice favored by each accumulator (right accumulator: right choice; left accumulator: left choice). As for the neural data, we used 10-fold cross-validation within each session, and the log odds of each trial were taken from the decoder trained on the other nine folds.

#### Across-trial correlation of neural or model DVs

We quantified the trial-to-trial correlations between the levels of left and right neural and model DVs at stimulus offset (taken as the average values between 0.9 to 1 s from stimulus onset). To factor out the effect of stimulus and choice on the correlations, we computed separate correlations for each stimulus category (right or left stronger according to the generative mean) and for each choice (right or left stronger), followed by averaging these four correlation coefficients. Correlations between model DVs were evaluated for two models: the best-fitting model, and a version of the latter without cross-inhibition (i.e., all parameters identical to the parameters estimated for the best-fitting model, except that FFI=0).

#### Within-trial correlation of neural or model DVs

To quantify the within-trial correlations between winning and losing neural (or model) DVs, we first z-scored the single-trial DV time series (for each selected trial epoch, see below) and detrended each DV. Detrending removed the shared ramping component shared by both DVs. We then computed the cross-correlograms of the resulting time courses for the lag range from -100 to + 100 ms, for the early stimulus interval (0 to 0.5 s from stimulus onset). For the neural DVs, this was also done for the pre-stimulus interval (-0.5 to 0 s from stimulus onset). Correlations between model DVs were again evaluated for the best-fitting model, and a version of the latter without cross-inhibition.

#### Contributions of winning and losing PMd/M1 neural DVs to confidence or RTs

We regressed the time courses of neural DVs from left and right PMd/M1 on confidence or RTs. For each subject, DV time courses were extracted from decoder outputs (log-odds) for each hemisphere. Neural DVs were averaged within sliding windows of 350 ms (step size: 100 ms) across the trial, producing time-resolved DV estimates. For each window and trial, we defined DV_left as “winning DV” and DV_right as “losing DV” for left choices, and conversely for right choices.. The design matrices of the models contained three regressors: evidence strength (absolute value of delta contrast) and the winning and losing neural DV. Trial-wise DV values, evidence strength and RTs were z-scored within session. This model was fit to single-trial RTs (linear regression) or to confidence reports (logistic regression). In addition to individual DV contributions, we computed an asymmetry index defined as the difference between the absolute regression weights of the winning and losing DVs (|β_win| − |β_lose|) for the time window centered at stimulus offset (-175 to 175 ms from stimulus offset), capturing the relative influence of the two competing accumulators.

#### Isolating components of association cortex DVs orthogonal to the PMd/M1 DV

Neural DV time courses were computed, again in units of log-odds, for multiple bilateral ROIs including PMd/M1. Neural DVs in association cortex tended to be strongly correlated with neural DVs in PMd/M1, which likely reflects a mixture of genuine neural correlations as well as spurious correlations (e.g., volume conduction, leakage of the spatial filters for source reconstruction). To estimate the unique contributions of neural DVs from all areas other than PMd/M1 to confidence, we used an established linear approach^89^ to first orthogonalize each of these DVs to the DV from bilateral PMd/M1 before fitting regression models on confidence. This was done to preclude strong correlations between regressors in multiple regression models. Specifically, we averaged the single-trial time courses of neural DVs from each area within sliding time windows (350 ms window, 100 ms step). From the resulting, smoothed time courses, we extracted across-trial vectors of local DVs, yielding as many such vectors as there were trial time windows. We refer to the across-trial vector of the PMd/M1 DV as “reference vector”. We then computed the linear projection of each DV-vector (i.e., from each association area) onto the reference vector, the latter normalized to unit length. This projection yielded a scalar weight (one per time point). We linearly removed from the DV-vectors of each association area the component shared with the reference vector, by subtracting the reference vector, scaled with the weight. The resulting “residual neural DVs” were now orthogonal to the PMd/M1 DV, ensuring that they captured unique decision-related activity in association cortex that was not explained by PMd/M1. We determined whether each residual neural DV supported the chosen action on a given trial by comparing the sign of the residual DV in each area with the sign of the subject’s choice. Positive alignment indicated activity favoring the selected option.

#### Contribution of winning and losing DVs in association cortex to confidence or RTs

For each trial and time point, we first defined winning residual neural DVs within each time window as the DVs whose sign matched the chosen option, and conversely for losing residual DVs, separately for each time window (see above). We took the absolute values of these DVs and separately averaged those values across winning DVs and losing DVs. We used regression models to test how trial-by-trial variations in the absolute value of winning and losing residual DVs related single-trial RTs or confidence reports. The design matrices of the models contained three regressors: the evidence strength (absolute value of delta contrast), the mean of the absolute values of residual DVs across winning areas, and the mean of the absolute values of residual DVs across all losing areas. Regressors and RTs were z-scored within session. The model was fit to single-trial RTs (linear regression) or to single-trial confidence reports (logistic regression), separately for each participant.

#### Selection of ROIs for analysis of neural DVs

From each of the 180 areas for which choice decoding was performed, we selected ROIs as candidate “neural DV” if their decoder outputs met the following three criteria (Table S1).

##### Choice information beyond PMd/M1

The decoder output from each region was averaged over the last 100 ms of stimulus presentation and the component shared with the PMd/M1 DV was removed by orthogonal projection (see above). The sign of the residual DV was then used to predict the participant’s choice, yielding prediction accuracy. We tested if the prediction accuracy was larger than chance (one-sided permutation test). P-values were corrected across areas using FDR correction (q < 0.05), separately for each criterion.

##### Evidence dependence of DV ramp

For each correct trial, we estimated the slope of the decoder output (linear fit over 200 to 1000 ms from stimulus onset). Pearson correlations between single-trial *ΔC* and DV ramping slopes were computed separately for both choices and averaged, giving one evidence-scaling value per subject. We tested at the subject-level for each area if the average Pearson correlation was larger than zero using a one-sided permutation test.

##### Increase of DV variance over time

For each session, and for left and right choices separately, we computed the across-trial variance of the decoder output at every time point and fitted a line to variance over time from 500 to 1000 ms. We tested at the participant-level for each area if the average slope was significantly larger than zero (one-sided permutation test).

#### Estimation of ramp onset of neural DVs in association cortex

For each participant, ROI, and candidate neural DV, we estimated the onset of the DV ramp relative to stimulus onset. Within each session, single-trial DV time courses were sign-flipped so that evidence for the chosen option was positive. DV time courses were then averaged across sessions. Each DV time course was normalized by its value at 1350 ms (around the peak of the DV time courses), and its local slope was computed in a time-variant (sliding window regression; 100 ms window length). The threshold for ramp onset detection was determined as the mean baseline slope (−500 to 0 ms) plus two standard deviations of that slope, which we used as proxy for the noise of our slope estimates. To compute the standard deviation of baseline slopes, trials were resampled with replacement 1000 times and baseline slopes were computed for each iteration and averaged across all 100 ms time windows within the baseline interval. A ramp onset was detected if the first threshold-crossing satisfied the following conditions: (i) the window start was at or after stimulus onset; (ii) subsequent slope estimates up to the DV peak were positive for at least 75% of time windows. Requiring the DV to keep rising until the decision excluded transient up-and-down fluctuations. The ramp onset was defined as the first time point of the 100 ms window at the detected threshold-crossing. ROIs in individual participants without such threshold-crossing were excluded from the latency analysis (range across ROIs: from 0 to 4 participants).

#### Statistical significance tests

To test differences of behavioral and neural measures against zero we used two-sided permutation tests (10,000 permutations). To account for multiple comparisons, we used cluster-based permutation tests for time-variant regression analyses and FDR correction with a threshold of q < 0.05 for across samples analyses. For a conservative assessment of significance for cortical maps of decoder accuracy, we assessed the local decoding accuracy with respect to the accuracy level of the ROI with the minimum accuracy level across the group-level map. To this end, the latter accuracy level (which may be above chance level due to volume conduction) was subtracted from each individual map of decoding accuracy values across ROIs. One-sample, two-sided permutation tests (10,000 permutations) were then performed for each region, testing whether the residual values were significantly greater than zero across subjects. Resulting p-values were FDR-corrected with a threshold of q < 0.01. For time-variant regression analyses, participant-wise regression coefficients were averaged and then tested against zero across participants using cluster-based permutation tests.

## Acknowledgements

We thank Karin Reimann for helping with subject recruitment; Marlene Petersson, Barbora Schwarzova, and Christiane Reissmann for helping with data collection. We thank Roozbeh Kiani and Xiao-Jing Wang for helpful discussion.

## Funding

Deutsche Forschungsgemeinschaft (DFG, German Research Foundation): SFB 936—178316478— A7 (T.H.D.) and Z3 (T.H.D.)

German Federal Ministry of Education and Research (BMBF): 01EW2007A (THD) Federal State of Hamburg consortium LFF-FV76 (THD)

ERA-NET NEURON consortium IMBALANCE (THD. and JdR)

## Author contributions

Conceptualization: AT, JdR, KT, THD

Methodology: AT, KT

Investigation: AT, AA

Visualization: AT, THD

Funding acquisition: THD

Project administration: THD

Supervision: THD

Writing – original draft: AT, THD

Writing – review & editing: AT, AA, JdR, KT, THD

## Data and code availability

Custom-written codes are available at https://github.com/DonnerLab/Multi-area-Decision-Dynamics-Across-Human-Cortex-Shape-Confidence.git. All data will be made available upon publication.

