## Supplementary information for "Multi-area Decision Dynamics Across Human Cortex Shape Confidence"

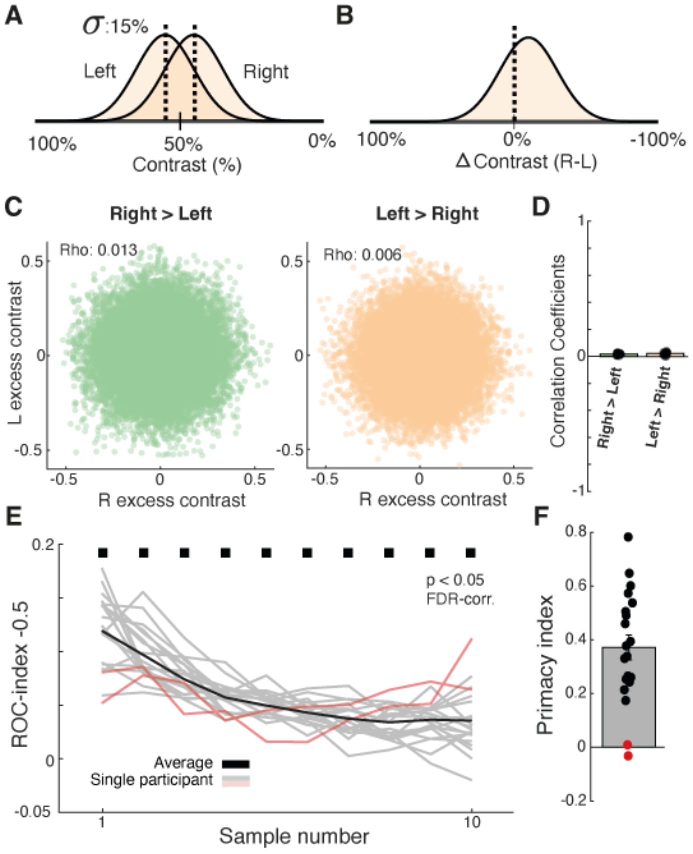


**Figure S1. Task design and behavior.** **(A)** Generative distributions of contrasts for the left and right stimulus stream in an example trial**.** **(B)** Generative distribution of delta-contrast for the same trial. **(C, D)** Demonstration that the sample contrast fluctuations on the left and right side are independent within each stimulus category defined by the generative mean. **(C)** Two-dimensional distributions of excess contrasts (difference of sample from generative mean) on left and right side, for an example participant. Distributions are plotted separately for each stimulus category. **(D)** Left and right sample correlations as in C, for the whole group. Bars, group average; vertical lines SEM. **(E)** Psychophysical kernel quantifying the impact of contrast difference fluctuations on choice as AUC of the ROC. Gray lines, individual participants; black line, group average; red lines, participants with a primacy index ≤ 0. Black horizontal squares, $p < 0.05$ (two-sided permutation-tests against 0, FDR-corrected). **(F)** Primacy index for all participants, computed as the difference between the summed ROC indices for the first and last five samples, normalized by the sum of their absolute values. Positive values indicate primacy (greater influence of early samples on choice), whereas negative values indicate recency (greater influence of late samples). Red dot, participant with a primacy index ≤ 0.

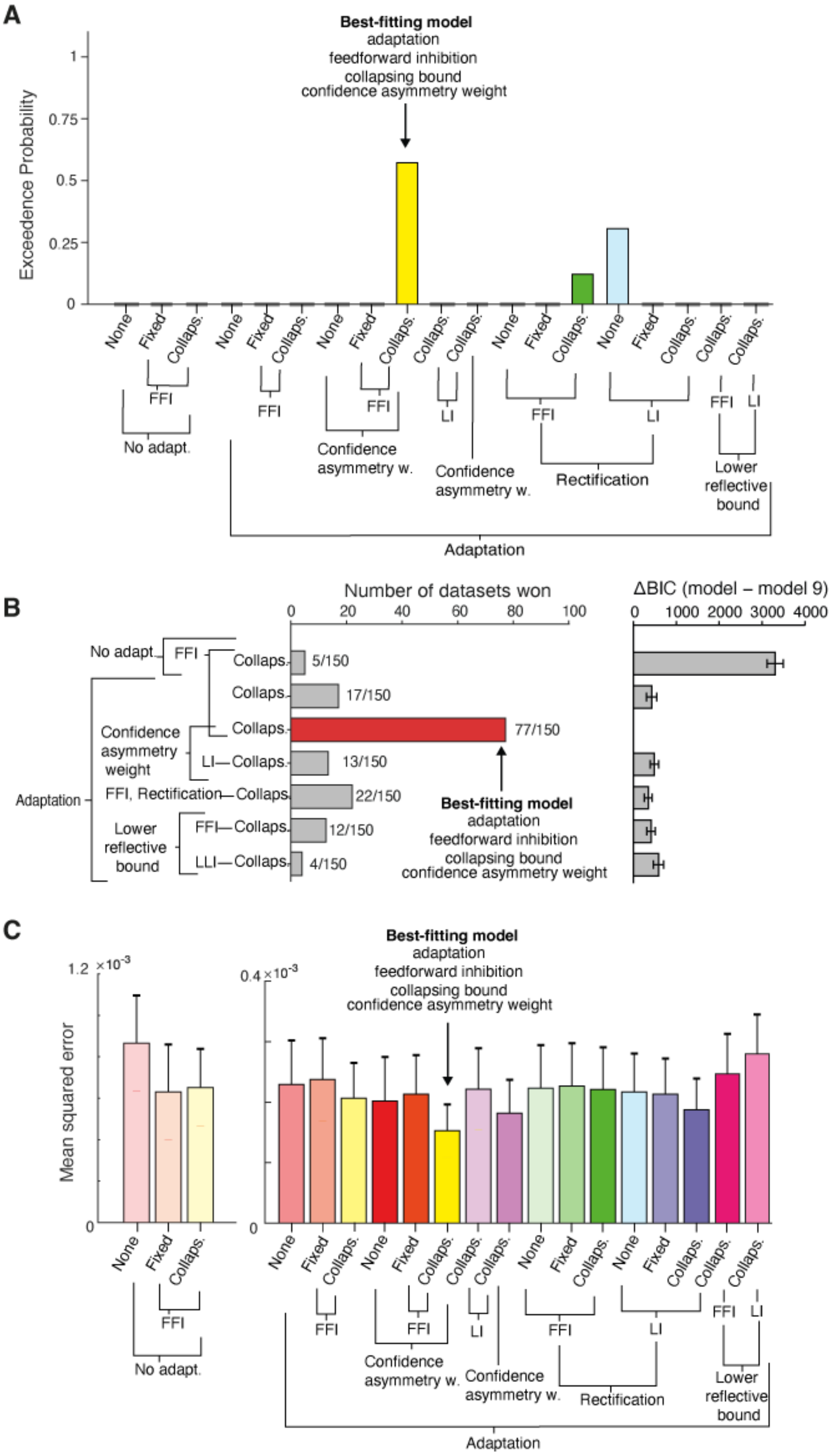


**Figure S2. Model selection. (A)** Model comparison. Bars show the probability that a given model best explains behavior across the population relative to all other models (exceedance probability), estimated by random-effects Bayesian model comparison. **(B)** Model recovery. Number of synthetic datasets, out of 150 generated by the best-fitting model (Model 9, see Methods), in which each of the seven candidate models had the lowest BIC. Model 9 was recovered in 77/150 datasets (51,3%; chance, 12.5%). Right panel: difference in BIC between each candidate model and the best-fitting model averaged across the 150 synthetic datasets. Bars, mean across datasets; error bars, SEM. **(C)** Best-fitting model also best captures the psychophysical kernels (main Figure 1). This is quantified in terms of mean squared error between data and model predictions for psychophysical kernels. All panels: bars, group average; error bars, SEM. For all metrics, Model 9 fit the data best and is therefore used throughout the main paper.

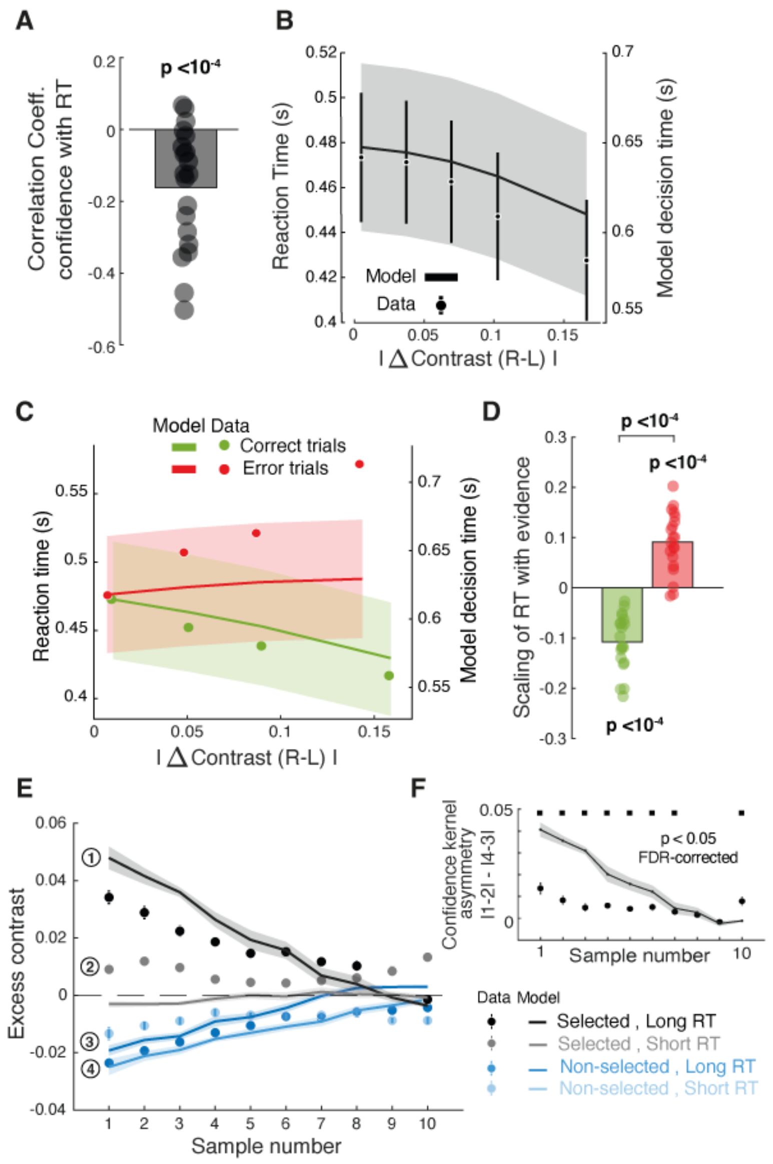


**Figure S3. Signatures of decision uncertainty in reaction time (RT). (A)** Correlation coefficients between confidence report and RT. Data points, individual participants; bars, group average. P-values obtained from two-sided permutation test against 0. **(B)** RT as a function of evidence strength (absolute value of delta contrast). **(C)** RT as a function of absolute value of delta contrast, split by correct and error trials. **(D)** Correlation between RT with evidence strength for correct (green) and error (red) trials. Data points, individual participants; bars, group average. P-values obtained from two-sided permutation test against 0. **(E)** Excess contrast fluctuations sorted by the side selected by the participant and short versus long reaction times (RT). **(F)** Asymmetry of confidence kernels, quantified as the difference between confidence kernels for selected (1-2) and non-selected (4-3) side in (E). Black horizontal squares indicate $p < 0.05$ (two-sided permutation-test against 0, FDR-corrected). Panels A,D: Bars, group average; data points, individual participants. All other panels: Data points, group average; vertical lines, SEM across participants; shaded area, SEM of model predictions.

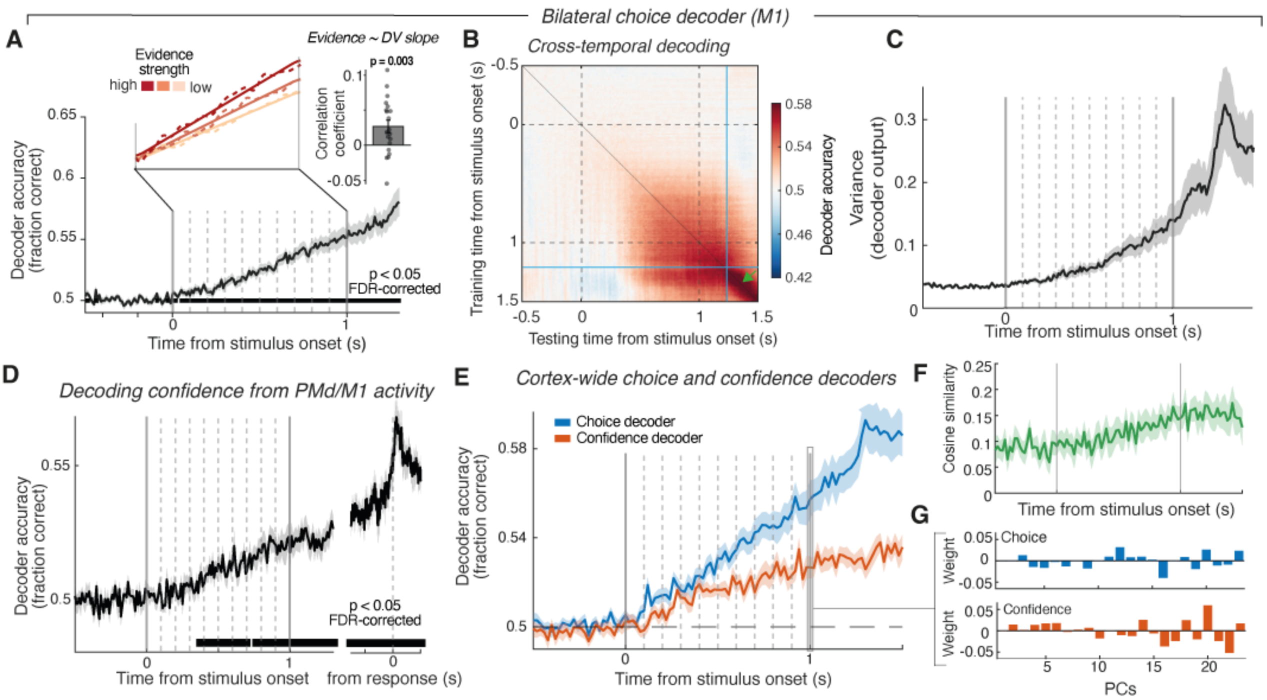


**Figure S4. Codes for integrated evidence and confidence in PMd and M1. (A)** Time-course of choice decoding from bilateral primary motor cortex (M1), aligned to stimulus onset and behavioral choice. Black line, group average, shaded area, SEM. Top left inset: decoder accuracy time-course for three levels of evidence strength. Dashed lines, data; solid lines: fitted slopes. Top right inset: single-trial correlation between the slope of decoder output and evidence strength (Methods). Ramping slopes were fitted for the interval 0.2 to 1 s from stimulus onset (Methods). Cross-validated decoder accuracy gradually ramped up during the 1s presentation of the evidence stream, followed by a transient peak aligned to the time of the behavioral choice report. **(B)** Temporal cross-decoding matrix of choice decoder from area M1 aligned to stimulus onset. Colored arrows indicate different part of the matrix reflecting different neural encoding of choice (see main text). The spatial pattern of choice-predictive activity was stable and gradually built up in amplitude during stimulus viewing, as reflected by the planar structure of the cross-temporal decoding matrix (i.e., similar accuracies for diagonal and off-diagonal elements)[1]. By contrast, during movement execution choice-related activity patterns were transient, with high decoding accuracy across the diagonal and small off-diagonal values (green arrow), in line with the transition between distinct neural activity subspaces for action planning and action execution found in animal work[2–4]. Blue line, group mean of RT. (**C**) Time-course of the variance of choice decoder output from bilateral primary motor cortex (M1), aligned to stimulus onset. Black line, group average, shaded area, SEM. **(D)** Time course of accuracy of confidence decoder trained and tested on bilateral PMd/M1 data. **(E)** Time course of accuracy of choice and confidence decoder trained on all bilateral cortical ROIs. **(F)** Time course of the cosine similarity between the weight vectors of the linear decoders trained to predict choice and confidence in E. **(G)** Decoder weights assigned to each PC by the choice (blue) and confidence decoder (orange) at stimulus offset.

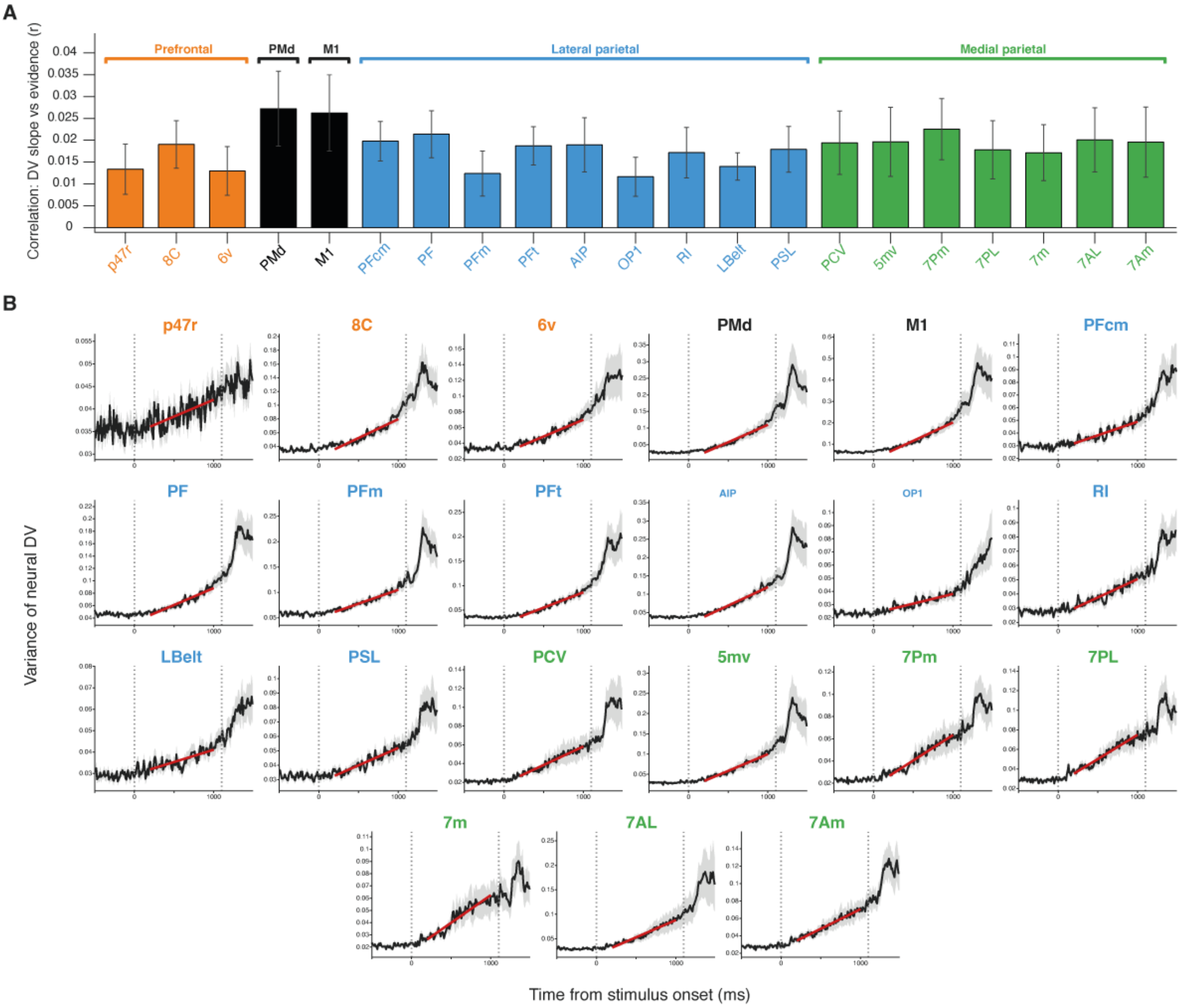


**Figure S5. Characterization of Neural DVs in association areas. (A)** Correlation coefficients of neural DVs slope of build-up activity during stimulus viewing with evidence strength. **(B)** Variance of neural DV over time for all selected areas. Variance had to increase linearly over time in the second half of stimulus presentation (in red).

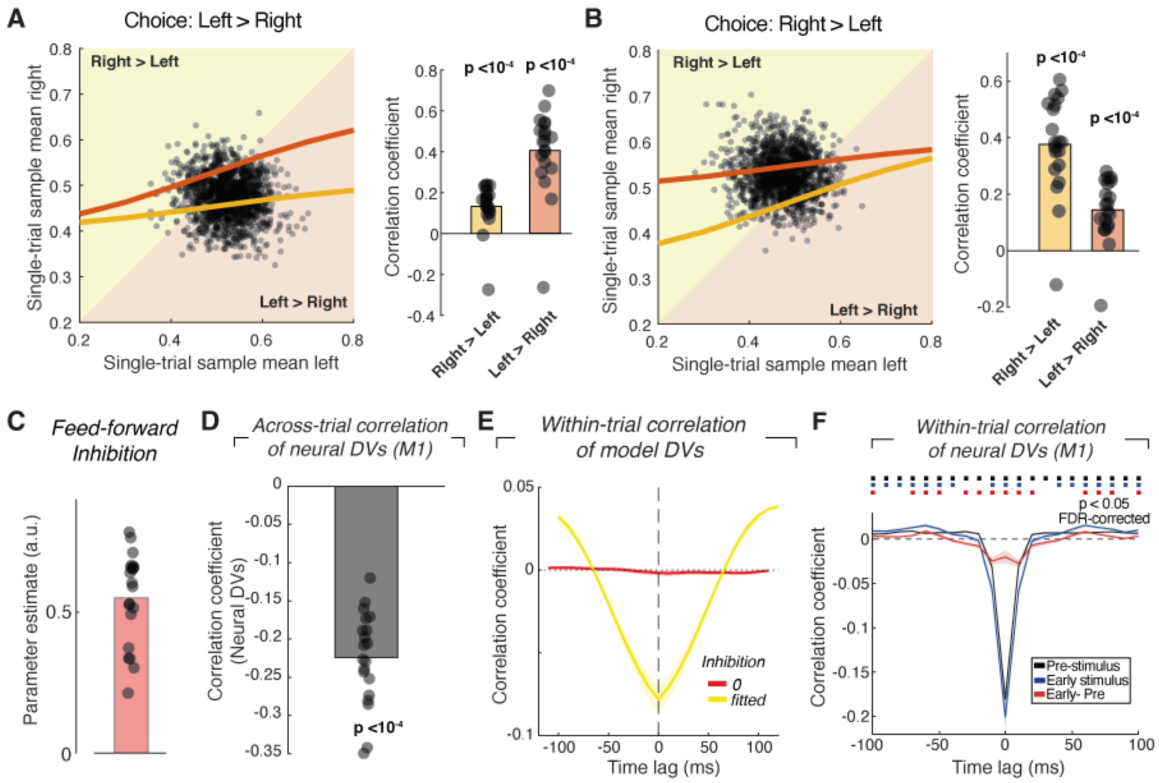


**Figure S6. Correlations between sample contrasts or winning and losing neural DVs. (A, B)** Across-trial correlations of average contrast levels (sample mean) presented on the right and left hemifield, after sorting by stimulus category and choice. **(A)** Left choice. **(B)** Right choice. Distributions of average contrast levels (sample mean) for example participant (same as Supplementary Figure 1C). Dots, single-trial sample means on left and right side. Note that the left and right average contrasts were, by design, negatively correlated across trials (rho = -0.27). However, when splitting trials by stimulus category, correlations were slightly positive (yellow and orange lines, rho = 0.22 and rho = 0.20 respectively). This reversal of the pooled correlation within strata is well-established in statistics [5]. It indicates that stimulus statistics cannot account for negative correlations between winning and losing neural DVs once stimulus category and choice are factored out. **(C)** Parameter estimates for inhibition. **(D)** Across-trial correlation between neural DV from motor cortex (M1) at the end of stimulus viewing (0.95 s after stimulus onset for neural DVs). Stimulus category and choice have been factored out (Methods). **(E)** Within-trial correlation between winning and losing model DVs in the early stimulus presentation (from 0 to 0.5 s from stimulus onset) for different level of cross-inhibition. Correlation is around 0 without inhibition between the two accumulators. **(F).** Within-trial correlation between winning and losing neural DVs from M1 time courses from two trial segments: pre-stimulus interval and early stimulus presentation (first 500 ms after stimulus onset) and their difference (in red), shown as cross-correlograms. Marks, $p < 0.05$ (FDR-corrected); p-values from permutation tests against 0 or between the two segments (in red)**.**

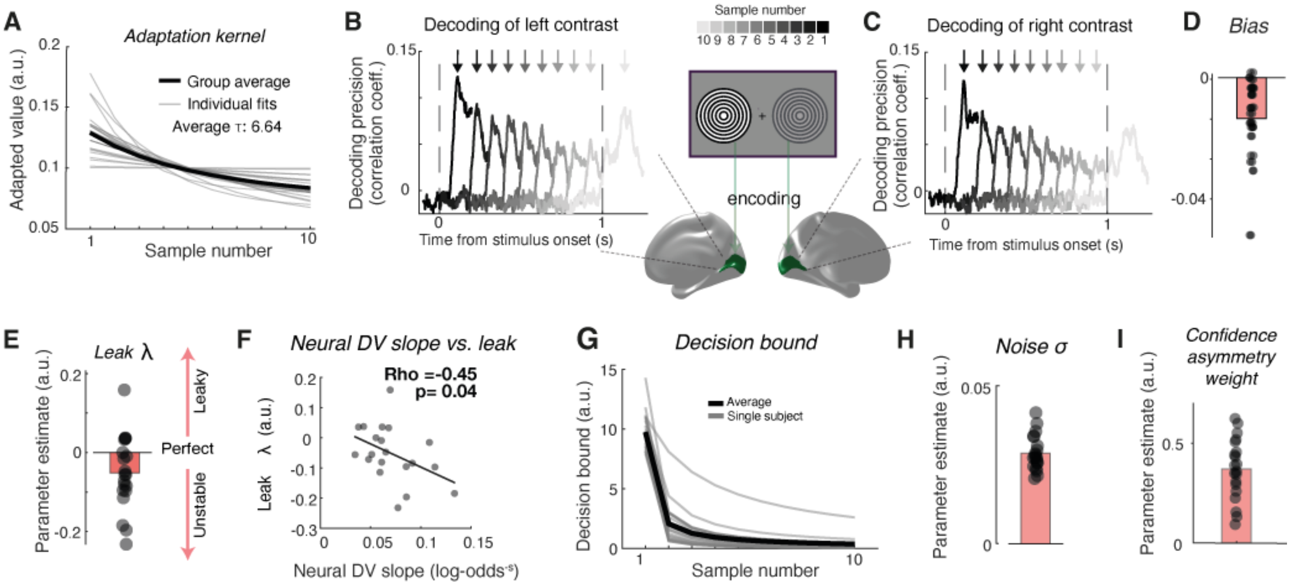


**Figure S7. Relationship between accumulator model and neural data.** **(A)** Estimated dynamics of adaptation. Gray lines, individual fits; black line, group average fit. All individual model fits exhibited a decaying adaptation function, in line with the V1 data shown in panels B,C. **(B)** Time courses of decoding of individual contrast samples in the left visual hemifield from right V1 response patterns. Decoding precision is expressed as cross-validated correlation between decoded and presented contrast (Methods). Lines group average time courses. Arrows, time point of peak decodability per sample position**. (C)** Same as B, but for contrast samples in right hemifield, decoded from left V1. The data show an attenuation of the encoding precision for contrast samples. **(D)** Parameter estimates for fixed bias. **(E)** Parameter estimates for leak. Parameters were negative in most participants, which is indicative of categorization dynamics[6]. **(F)** Pearson correlation between individual leak parameter and PMd/M1 choice decoder slopes. Leak parameters correlated with the slope of the neural DV in PMd/M1 (bilateral choice decoder). This is consistent with the idea that subjects that tended to commit to a choice early during stimulus viewing, were captured by a more negative leak and faster build-up of the neural DV. **(G)** Estimated dynamics of decision bound. Gray lines, individual fits; black line, group average fit. **(H-J)** Parameter estimates for (H) noise and (I) confidence asymmetry weight. Data points, individual participant estimates; bars, group average.

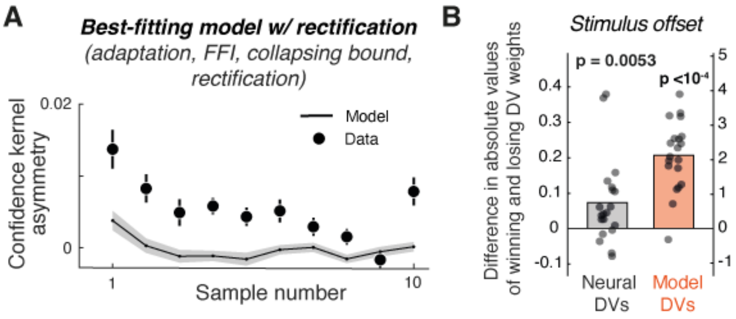


**Figure S8. Predictions for best fitting model with rectification of the DVs. (A)** Asymmetry of confidence kernels quantified as in Figure 1G. Data points, group average; vertical lines, SEM across participants; Shaded error bar, model prediction. **(B)** Difference between absolute value of beta weights for winning and losing neural and model DVs for model with rectification at the end of stimulus viewing (1 s after stimulus onset). "Neural DV" plots same data as in Figure 3D for comparison. Effects are still significant after excluding the two participants without a primacy effect in the psychophysical kernels ($p = 0.0002$ for Neural DVs and $p = 0.008$ for Model DVs two-sided permutation test against zero). Bars, group average; data points, individual participants.

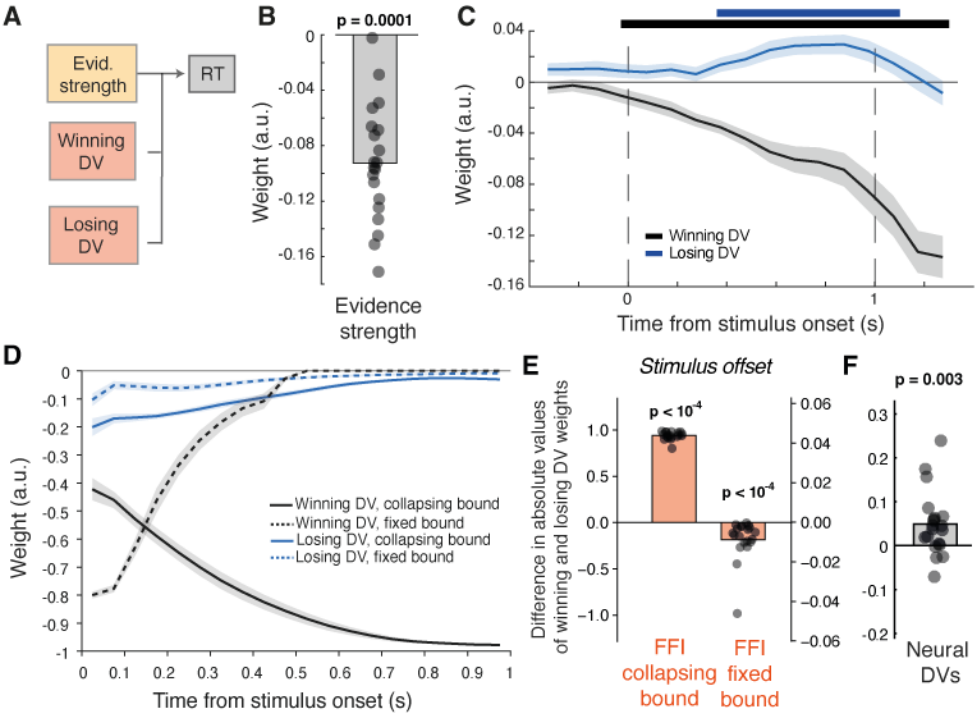


**Figure S9. Relationship between neural and model DVs and RT. (A)** Schematic of linear regression model predicting single-trial reaction times from evidence strength and level of neural DVs over time (350 ms moving window, 100 ms steps, see Methods). **(B)** Beta coefficients for evidence strength at the end of stimulus viewing (1 s after stimulus onset). **(C)** Beta weights over time for winning versus losing neural DVs. Marks, $p < 0.05$ (cluster-based permutation test)**.** Lines, group average; shaded areas, SEM. **(D)** Beta weights over time for the winning (black) and losing (blue) model DV of the best-fitting model (solid lines) and of the same model with a fixed bound (dashed lines), from a regression of evidence strength and the winning and losing DV on model bound crossing time. With a fixed bound, once the winning DV has crossed the bound on all trials, the winning DV remains at a constant value equal to the bound, so its weight on model decision time becomes zero. With a collapsing bound, the weight of the winning DV becomes more negative over time, as more trials cross the bound, while the beta of losing DV goes to zero. Lines, group average; shaded areas, SEM across participants. **(E)** Difference between the absolute values of the weights of the winning and losing model DVs at the end of stimulus viewing for the two model architectures. **(F)** Same as (E) for the neural DVs in PMd. In all panels, bars are group average, data points are individual participants and p-values are from two-sided permutation test against zero.

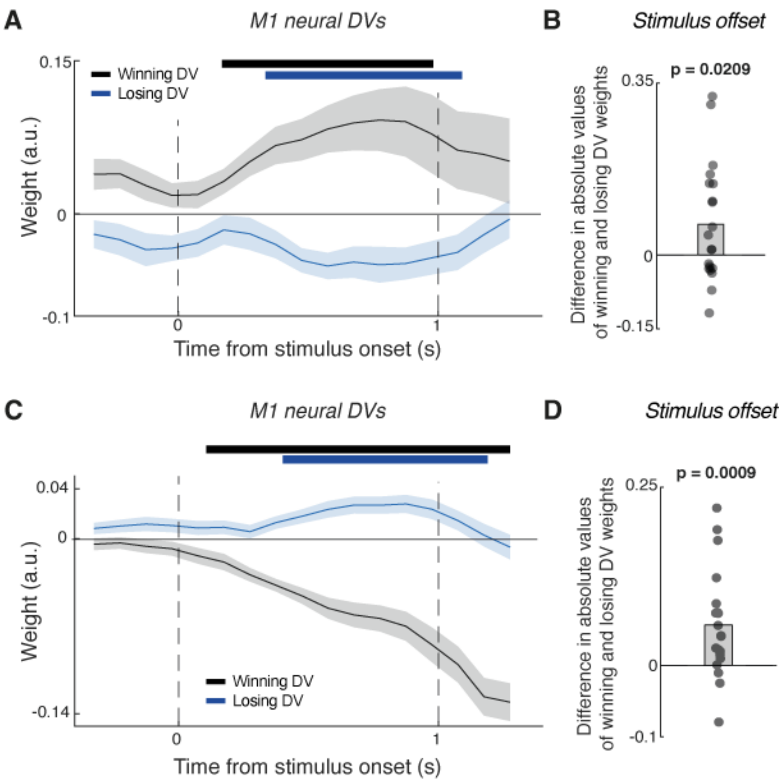


**Figure S10. Unique contributions of competing neural DVs in M1 to confidence and RT. (A-B)** Unique contributions of competing neural DVs in M1 to confidence. **(A)** Time course of beta weights for winning versus losing neural DVs. The latter were evaluated on a sliding-window basis (350 ms window length; Methods. Marks, $p < 0.05$ (cluster-based permutation test). Lines, group average; shaded areas, SEM. **(B)** Difference between absolute value of beta weights for winning and losing neural DVs at the end of stimulus viewing (1 s after stimulus onset). **(C-D)** Unique contributions of competing neural DVs in M1 to RTs. **(C)** Beta weights over time for winning versus losing neural DVs. Marks, $p < 0.05$ (cluster-based permutation test)**.** Lines, group average; shaded areas, SEM. **(D)** Difference between absolute value of beta weights for winning and losing neural DVs at the end of stimulus viewing (1 s after stimulus onset). P-values in all panels are from two-sided permutation tests against 0. Bars, group average; data points, individual participants.

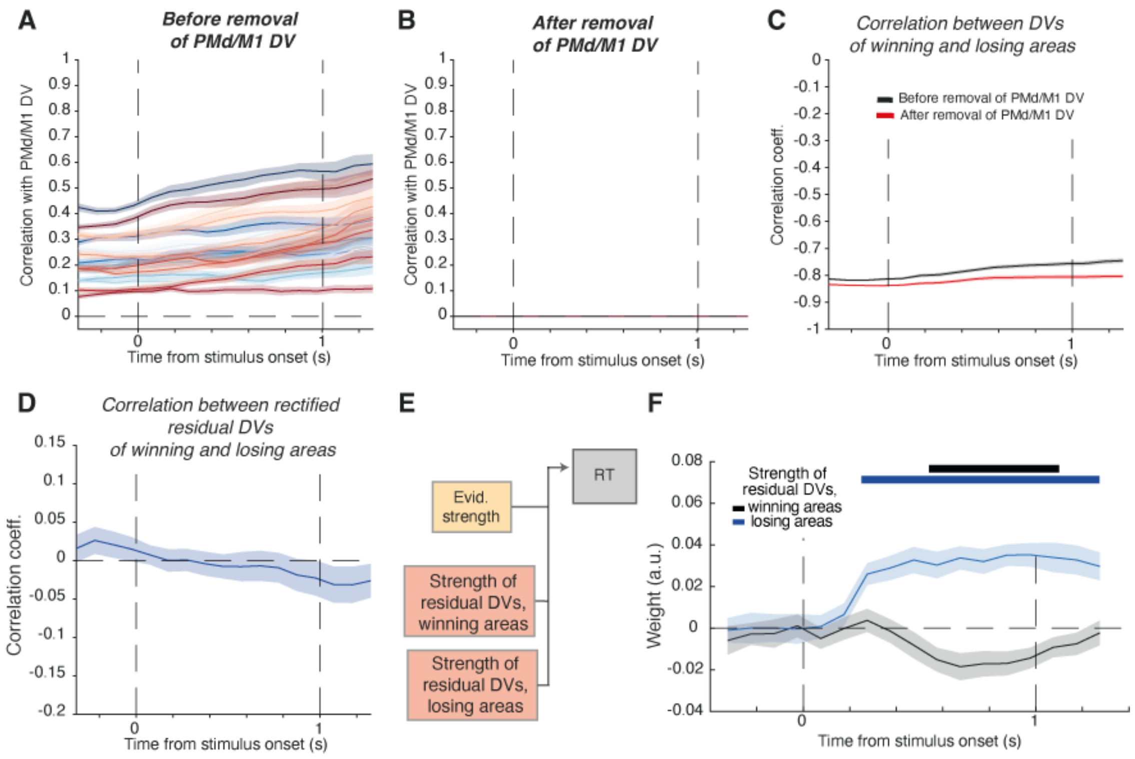


**Figure S11. Correlations between neural DVs in association cortex, PMd/M1 DV, and RT. (A-B)** Time course of correlation between neural DVs within each of the 22 selected ROIs from main Figure 4A with the PMd/M1 DV, before (A) and after (B) removal of each area's component shared with the PMd/M1 (Methods). **(C)** Correlation between the mean of winning residual DVs and mean of losing residual DVs, computed before (black line) and after (red line) removal of the PMd/M1 DV. **(D)** Correlation between the absolute values of mean of winning residual DVs and mean of losing residual DVs, computed after removal of the PMd/M1 DV. **(E)** Schematic of linear regression model predicting single-trial reaction times from evidence strength and the strength of residual DVs from losing and winning areas after PMd/M1 DV has been projected out (see Methods). **(F)** Evolution over time (350 ms moving time window, 100 ms steps) of regression weights linking RT to the strength of losing and winning areas residual DVs. Horizontal bars, $p < 0.05$ (cluster-based permutation test)**.** Lines, group average; shaded areas, SEM.

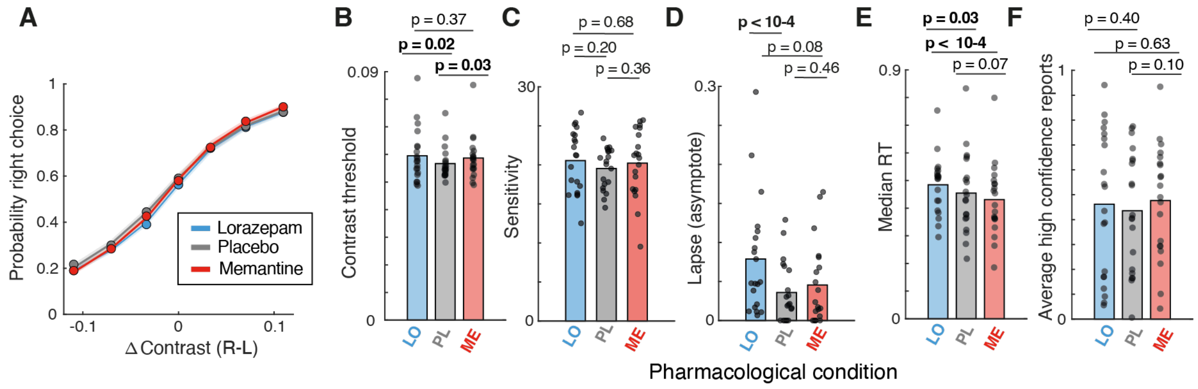


**Figure S12. Subtle effects of pharmacological interventions on task behavior.** Quantification of choice behavior split by pharmacological condition. **(A)** Psychometric functions quantified as probability of right-side choice as a function of average delta contrast. Data points, group average; lines, psychometric function fits. **(B)** Average delta-contrast threshold per participant and condition. **(C)** Sensitivity. **(D)** "Lapse" parameter quantifying the distance of the two fitted asymptotes from 0 or 1, respectively. **(E)** Median RT. **(F)** Fraction of high-confidence reports. In B-F, bars are group average and data points are participants. Overall drug effects were subtle, reflecting the low dosages applied, and there was no statistically significant effect on the fraction of high-confidence reports. P-values obtained from two-sided permutation test.

| ROI | Area name | Residual accuracy (%) [95% CI] | Variance slope ($s^{-1}$) [95% CI] | Evidence scaling r [95% CI] |
|---|---|---|---|---|
| **Reference areas (PMd / M1)** | | | | |
| **6a (PMd)** | Area 6 anterior | | 0.1239 [0.053, 0.1942] | 0.021 [0.0052, 0.037] |
| **6d (PMd)** | Dorsal area 6 | | 0.1581 [0.0565, 0.2598] | 0.0296 [0.013, 0.046] |
| **M1 (4)** | Primary motor cortex | | 0.2040 [0.0978, 0.3102] | 0.027 [0.011, 0.044] |
| **Prefrontal areas** | | | | |
| **p47r** | posterior 47 rostral (lateral frontal lobe) | 50.6 [50.0, 51.2] | 0.0092 [0.0014, 0.0169] | 0.015 [0.004, 0.026] |
| **8C** | Area 8C (lateral frontal lobe) | 50.8 [50.2, 51.3] | 0.0728 [0.0362, 0.1094] | 0.019 [0.009, 0.029] |
| **6v** | Ventral area 6 | 51.3 [50.5, 52.2] | 0.0554 [0.0274, 0.0834] | 0.013 [0.004, 0.024] |
| **Lateral parietal areas** | | | | |
| **PFcm** | Area PF caudomedial (inferior parietal lobule) | 51.2 [50.5, 51.8] | 0.0267 [0.0082, 0.0452] | 0.019 [0.013, 0.028] |
| **PFt** | Area PF tenuicorticalis | 52.0 [51.3, 52.8] | 0.0716 [0.0373, 0.1059] | 0.02 [0.011, 0.029] |
| **PF** | Area PF complex (inferior parietal lobule) | 52.0 [51.3, 52.8] | 0.0686 [0.0363, 0.1009] | 0.024 [0.017, 0.033] |
| **PFm** | Area PF medial (inferior parietal lobule) | 52.3 [51.4, 53.1] | 0.0618 [0.0216, 0.1020] | 0.020 [0.000, 0.022] |
| **AIP** | Anterior intraparietal area | 52.4 [51.5, 53.3] | 0.1217 [0.0614, 0.1821] | 0.020 [0.007, 0.033] |
| **OP1** | Operculum parietale 1 (SII) | 50.6 [50.0, 51.2] | 0.0160 [0.0031, 0.0288] | 0.013 [0.005, 0.02] |
| **RI** | Retroinsular cortex | 50.9 [50.4, 51.4] | 0.0355 [0.0170, 0.0539] | 0.017 [0.005, 0.028] |
| **LBelt** | Lateral belt complex | 50.7 [50.3, 51.2] | 0.0201 [0.0069, 0.0333] | 0.014 [0.007, 0.020] |
| **PSL** | PeriSylvian language area | 51.4 [50.8, 52.1] | 0.0299 [0.0090, 0.0509] | 0.019 [0.008, 0.003] |
| **Medial and superior parietal areas** | | | | |
| **PCV** | Precuneus visual area | 51.6 [50.6, 52.6] | 0.0378 [0.0080, 0.0677] | 0.017 [0.002, 0.031] |
| **5mv** | Medial-ventral area 5 (Superior parietal lobule) | 51.3 [50.5, 52.1] | 0.0927 [0.0229, 0.1624] | 0.02 [0.004, 0.035] |
| **7Pm** | Posterior-medial area 7 (Superior parietal lobule) | 52.0 [51.0, 53.1] | 0.0409 [0.0107, 0.0711] | 0.015 [0.002, 0.028] |
| **7m** | Medial area 7 (Superior parietal lobule) | 52.4 [51.2, 53.7] | 0.0314 [0.0036, 0.0593] | 0.018 [0.002, 0.035] |
| **7Am** | Medial area 7a (Superior parietal lobule) | 52.0 [51.1, 53.0] | 0.0480 [0.02, 0.0760] | 0.016 [0.002, 0.030] |
| **7PL** | Posterior-lateral area 7 (Superior parietal lobule) | 52.7 [51.8, 53.7] | 0.0474 [0.0146, 0.0802] | 0.018 [0.0035, 0.034] |
| **7AL** | Lateral area 7a (Superior parietal lobule) | 52.2 [51.0, 53.4] | 0.0796 [0.0136, 0.1457] | 0.015 [0.001, 0.030] |

**Table S1.** Neural decision-variable (DV) statistics for the selected ROIs. Values are means across subjects (N = 20) with 95% confidence intervals.
